# Equivalent noise characterization of human lightness constancy

**DOI:** 10.1101/2021.06.04.447171

**Authors:** Vijay Singh, Johannes Burge, David H. Brainard

## Abstract

A goal of visual perception is to provide stable representations of task-relevant scene properties (e.g. object reflectance) despite variation in task-irrelevant scene properties (e.g. illumination, reflectance of other nearby objects). To study such stability in the context of the perceptual representation of lightness, we introduce a threshold-based psychophysical paradigm. We measure how thresholds for discriminating the achromatic reflectance of a target object (task-relevant property) in rendered naturalistic scenes are impacted by variation in the reflectance functions of background objects (task-irrelevant property), using a two-alternative forced-choice paradigm in which the reflectance of the background objects is randomized across the two intervals of each trial. We control the amount of background reflectance variation by manipulating a statistical model of naturally-occurring surface reflectances. For low background object reflectance variation, discrimination thresholds were nearly constant, indicating that observers’ internal noise determines threshold in this regime. As background object reflectance variation increases, its effects start to dominate performance. A model based on signal detection theory allows us to express the effects of task-irrelevant variation in terms of the equivalent noise, that is relative to the intrinsic precision of the task-relevant perceptual representation. The results indicate that although naturally-occurring background object reflectance variation does intrude on the perceptual representation of target object lightness, the effect is modest - within a factor of two of the equivalent noise level set by internal noise.

## INTRODUCTION

To support effective action, vision provides stable perceptual representations of the distal properties of objects. The computations that give rise to these representations start with the information in the proximal stimuli that are encoded by the retinas. These proximal stimuli depend on the intrinsic properties of the objects in the scene, on object-extrinsic properties of the scene (e.g. illumination), and on the observer’s particular viewpoint. A challenge for the visual system is to recover stable perceptual correlates of object-intrinsic properties across variation in other scene variables. Understanding the degree to which the visual system rises to this challenge, and how it does so, is an important goal of vision science (Helmholtz, 1896; Knill & Richards, 1996; Brascamp & Shevell, 2021; Geisler, 2008; Burge, 2020; Wandell & Brainard, in press).

Here we consider the perceptual task of representing the reflectance of an object embedded in a scene, based on the light reflected to the eye from the object and the rest of the scene. The perceptual correlate of object surface reflectance is its perceived color or, in the special case of achromatic objects, its lightness. Computing a stable color or lightness representation poses a challenge to the visual system because the retinal image of the object varies with the object’s reflectance, the spectral irradiance of the illumination, the position and pose of the object in the scene, and the properties of other objects in the scene. The degree to which the visual system succeeds at stabilizing its color and lightness representations of objects, in the face of variation extrinsic to their reflectance, determines the degree to which the visual system achieves color and lightness constancy.

Under many circumstances, the visual system achieves a high degree of color and lightness constancy (Foster, 2011). Several theoretical frameworks have been developed to account for this ability. The frameworks attempt to explain how different cues are processed to form stable perceptual representations of object reflectance (Adelson, 2000; Smithson, 2005; Gilchrist, 2006; Kingdom, 2011; Foster, 2011; Brainard & Maloney, 2011; Brainard & Radonjić, 2014; Witzel & Gegenfurtner, 2018; Hurlbert, 2019; Murray, 2021). The underlying computations have been explained in terms of mechanistic gain control (e.g., von Kries, 1905; Whittle & Challands, 1969; Webster & Mollon, 1995; Land & McCann, 1971; Horn, 1974), cue combination (e.g., Maloney & Yang, 2001; Yang & Maloney, 2001), Bayesian inference (e.g., Brainard & Freeman, 1997; Brainard et al., 2006; Barron & Malik, 2012a; Allred & Brainard, 2013; Murray, 2020; see also Boyaci, Maloney, & Hersh, 2003; Bloj et al., 2004; Brainard & Maloney, 2011), learned computations (e.g., Singh, Cottaris, Heasly, Brainard, & Burge, 2018; Flachot & Gegenfurtner, 2018; Flachot & Gegenfurtner, 2021; Afifi, Barron, LeGendre, Tsai, & Bleibel, 2021), and application of principles of perceptual organization (Gilchrist, 2006; Adelson, 1993).

Color constancy and lightness constancy have been elucidated primarily with an experimental approach in which observers report on suprathreshold aspects of the color or lightness of a target object, across changes extrinsic to the target object’s reflectance.^1^ In these experiments, the target object’s reflectance is the task-relevant scene variable, while other aspects of the scene are task irrelevant. Observers’ reports are solicited using a variety of methods, including matching (e.g., Burnham, Evans, & Newhall, 1952; Gilchrist, 1977; Arend & Reeves, 1986; Brainard, Brunt, & Speigle, 1997), naming (e.g., Helson & Jeffers, 1940; Olkkonen, Witzel, Hansen, & Gegenfurtner, 2010), scaling (e.g., Schultz, Doerschner, & Maloney, 2006), and nulling (e.g., Helson & Michels, 1948; Jameson & Hurvich, 1955; Chichilnisky & Wandell, 1997; Brainard, 1998).

In the study of perception, discrimination experiments complement experiments that rely on suprathreshold reports. In a typical discrimination experiment, observers choose which of two stimuli has a larger physical value along some stimulus dimension. The stimulus difference is titrated to determine the smallest change that supports criterion discrimination performance. This smallest change is defined as threshold. For example, observers might be tasked with reporting which of two objects has a larger lightness value, in an effort to determine the human ability to discriminate different object surface reflectances. Mature theory links discrimination thresholds to the precision of the underlying perceptual representation (Green, 1966). Theory also exists for linking thresholds to properties of neural responses (Brindley, 1960; Green, 1966; Teller, 1984; Parker & Newsome, 1998).

Theory is less well developed for how to use discrimination experiments to address questions about perceptual constancy. In the case of color constancy, one approach is to measure observers’ ability to discriminate changes in scene illumination (Pearce, Crichton, Mackiewicz, Finlayson, & Hurlbert, 2014; Radonjić et al., 2016; Radonjić et al., 2018; Aston, Radonjić, Brainard, & Hurlbert, 2019; Alvaro, Linhares, Moreira, Lillo, & Nascimento, 2017), rather than to measure the ability to detect a change in object surface reflectance per se (for work that measures reflectance discrimination thresholds see Morimoto & Smithson, 2018). The idea is that if illumination changes are subthreshold, then the perceptual representations of both surface reflectance and illumination are stable across those illumination changes. However, it is unclear how the results of these experiments connect to and inform us about the stability of perceptual judgments across the larger illumination changes that occur in natural viewing (but see Weiss, Witzel, & Gegenfurtner, 2017). Another approach is to link discrimination thresholds to suprathreshold reports of perceived stimulus properties, an approach which has its origins in Fechner’s pioneering interpretation of Weber’s Law (Fechner, 1860). The idea is that both threshold and suprathreshold percepts are mediated by a common stimulus-response function whose properties depend on, and change with, viewing context. Although positing a common stimulus-response function holds promise (Nachmias & Sansbury, 1974; Hillis & Brainard, 2005; Hillis & Brainard, 2007b), there are cases in which the discrimination thresholds do not predict suprathreshold measures of lightness constancy made using well-matched stimuli (Hillis & Brainard, 2007a).

Here, we introduce a new approach to using discrimination experiments to study perceptual constancy. The approach is based on measuring how discrimination thresholds for a task-relevant scene property are affected by variation in a task-irrelevant scene property. The approach is conceptually similar to studying how contrast thresholds are affected by addition of random, unpredictable stimulus variation, usually introduced in the form of spatially white or pink contrast noise (Legge, Kersten, & Burgess, 1987; Pelli, 1990; Pelli & Farell, 1999). It is conceptually distinct in that the random, unpredictable variation is introduced in the distal scene properties (for related recent work, see Zhu, Yuille, & Kersten, 2021). We apply this approach to the study of lightness constancy in naturalistic scenes. First, we measure human ability to discriminate the achromatic surface reflectance of a target object in the absence of any target object-extrinsic variation. Next, we measure how these lightness discrimination thresholds change with the introduction of target object-extrinsic variation. Specifically, we introduce random, unpredictable variation to the background objects in the scene by varying their reflectance spectra—loosely, their colors (Lotto & Purves, 1999; Brown & MacLeod, 1997). The lightness discrimination threshold at each level of background object reflectance variation measures how difficult the lightness discrimination task is for that level of variation. The change in difficulty from baseline (i.e., no background object reflectance variation) quantifies the degree to which the background variation intrudes on the perceptual representation of target lightness.

As the variation in background object reflectances is increased, we find that discrimination thresholds are initially constant and then increase. To interpret these findings, we develop a model based on signal detection theory; the model is similar to those used to understand the effect of contrast noise on contrast thresholds (Legge, Kersten, & Burgess, 1987; Pelli, 1990). The model relates thresholds for the task-relevant variable (here, target object reflectance) to the amount of variation in the task-irrelevant variable (here, background object reflectance). The model allows us to express the effect of task-irrelevant variation in terms of equivalent noise. Equivalent noise is the amount of external task-irrelevant variation whose effect on the perceptual representation is the same as that of internal noise. We find that the intrusion of naturally occurring variation in background object reflectances on the perceptual representation of lightness is within a factor of two of the equivalent noise.

The paper is organized as follows: Section 2 (Methods) provides the experimental methods. Section 3 (Model) introduces the model used to interpret the data, and discusses the concept of equivalent noise in more detail. Section 4 (Results) reports the experimental results in the context of the model. Section 5 (Discussion) provides a summary. The Appendix describes a control experiment and provides supplemental figures and tables. Additional supplementary information is available online as indicated in Methods: Code and Data Availability.

## 2 EXPERIMENTAL METHODS

### Overview

We studied the effect of variability in object-extrinsic properties on the human ability to discriminate an object-intrinsic property. Specifically, we measured how variation in the reflectance spectra of background objects affects lightness discrimination thresholds, that is thresholds for discriminating object achromatic reflectance.^2^ We used a two-alternative forced-choice (2AFC) procedure (Figure 1). On each trial, observers viewed a standard image and comparison image, sequentially presented on a calibrated monitor for 250ms each. The inter-stimulus interval was 250ms (Figure 1a). The images were computer graphics renderings of 3D scenes. Each scene contained a spherical target object that appeared achromatic. The observers’ task was to report the image in which the target object was lighter. Across trials, we varied the luminous reflectance factor (LRF; American Society for Testing and Materials, 2017) of the target object in the comparison image while keeping the LRF of the target object in the standard image fixed. The LRF is the ratio of the luminance of a surface under a reference illuminant (here, the CIE D65 reference illuminant) to the luminance of the reference illuminant itself. The target object LRF was varied by scaling the surface reflectance spectrum of the target object, without changing its shape.^3^ The temporal order in which the standard and comparison images were presented was randomized on each trial.

**Figure 1:**
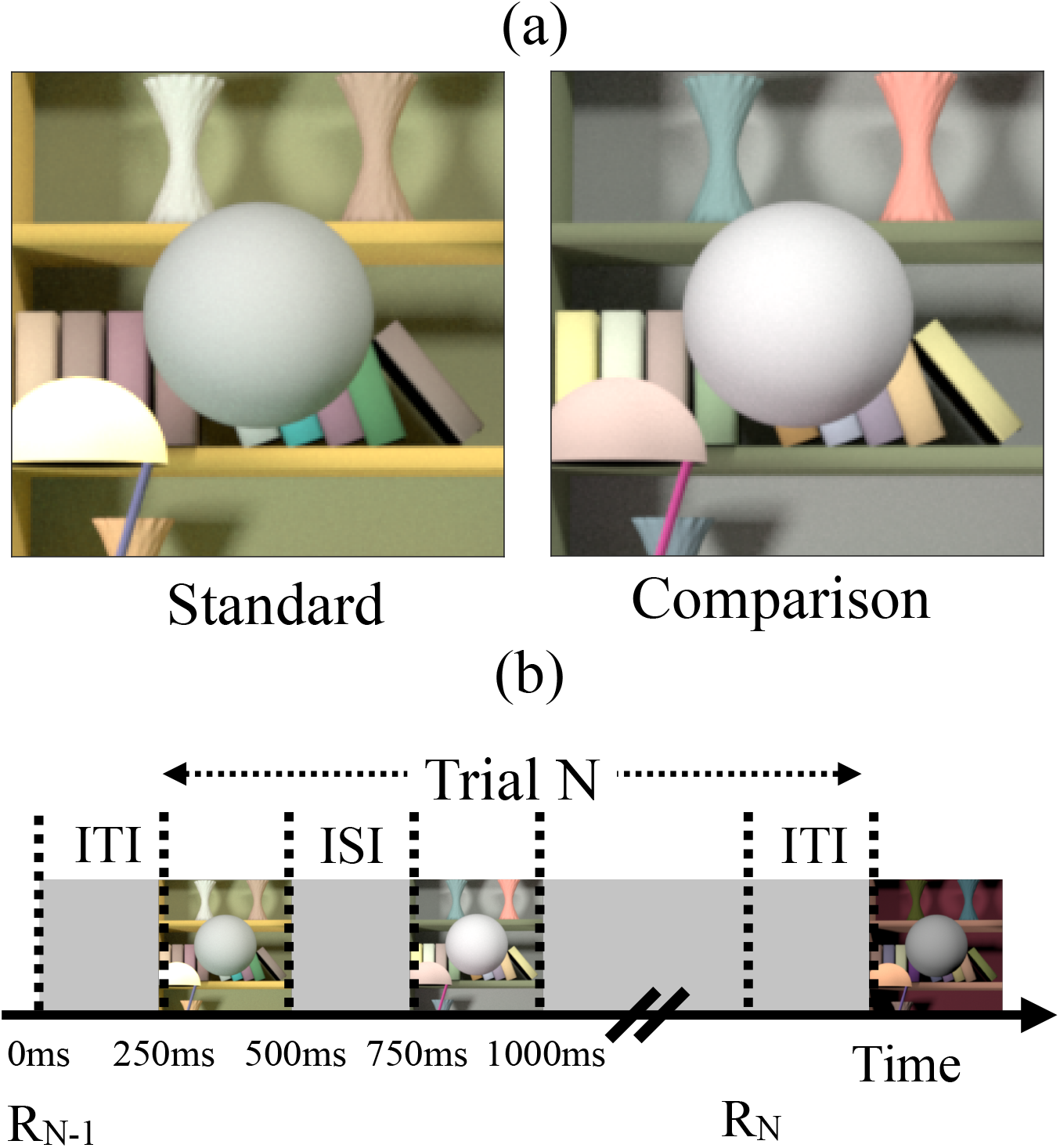
Psychophysical task. **(a)** On every trial of the experiment, human observers viewed two images in sequence, a standard image and a comparison image and indicated the one in which the spherical target object in the center of the image was lighter. Example standard and comparison images are shown. The images were computer graphics simulations. The simulated reflectance functions of the target were spectrally flat, and the spheres appeared gray. The overall reflectance of the target was held fixed in the standard images and differed between standard and comparison. Performance (proportion correct) was measured as a function of this difference to determine discrimination threshold. The reflectance spectra of objects in the background could be held fixed or vary between standard and comparison on each trial (as illustrated here). The order of presentation of the standard and comparison images was randomized from trial to trial. Discrimination thresholds were measured as function of the amount of variation in background object reflectances. **(b)** Trial sequence. R_N-1_ indicates the time of the observer’s response for the (N-1)^th^ trial. The N^th^ trial begins 250ms after that response (Inter Trial Interval, ITI). The N^th^ trial consists of two 250ms stimulus presentation intervals with a 250ms inter-stimulus interval (ISI). The observer responds by pressing a button on a gamepad after the second stimulus has been shown. The observer can take as long as he or she wishes before making the response, with an example response time denoted by R_N_ in the figure. The next trial begins 250ms after the response.

We recorded the proportion of times observers chose the comparison image as having the lighter target object at 11 values of the target object LRF. Figure 2 shows a psychometric function from a typical human observer. The proportion-comparison-chosen data were fit with a cumulative normal using maximum likelihood methods (see Methods: Psychometric Function). Threshold was defined as the difference between the LRF of the target object at proportion comparison chosen 0.76 and 0.50 (i.e., d-prime = 1.0 in a two-interval task), as determined from the cumulative normal fit.

**Figure 2:**
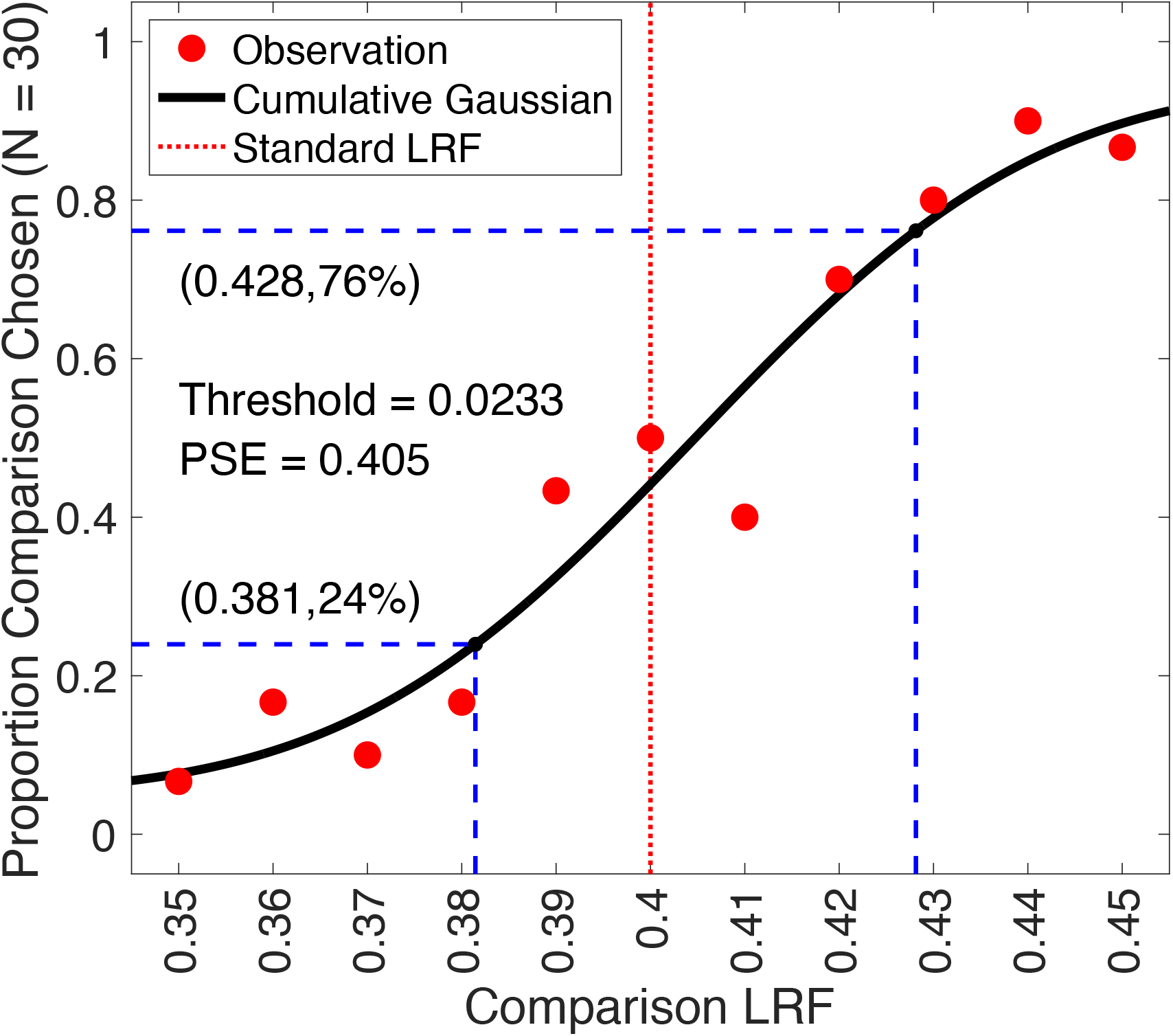
Psychometric function. We recorded the proportion of times the observer chose the target in the comparison image to be lighter, as a function of the comparison LRF. The LRF of the target object in the standard image was fixed at 0.4. The LRF of the target object in the comparison image were chosen from 11 linearly spaced values in the range [0.35, 0.45]. In each block, thirty trials were presented at each comparison LRF value. We fit a cumulative normal distribution to the proportion comparison chosen data using maximum likelihood methods. The guess and lapse rates were constrained to be equal and were restricted to be in the range [0, 0.05]. The threshold was measured as the difference between the LRF at proportion comparison chosen equal to 0.76 and 0.5, as predicted by the cumulative normal fit. This figure shows the data for Observer 2 for scale factor 0.00, for the block run in the first experimental session for that observer. The point of subjective equality (PSE, the LRF corresponding to proportion chosen 0.5) was close to 0.4 as expected and the threshold was 0.0233. The lapse rate for this fit was 0.05.

We measured lightness discrimination thresholds as a function of the amount of variability in the surface reflectances of the background objects in the rendered scenes. The reflectances of the background objects were chosen from a distribution of natural reflectances. The amount of variability was controlled parametrically by multiplying the covariance matrix of the distribution by a scalar (see Methods: Reflectance and Illumination Spectra). We measured thresholds for six logarithmically spaced values of this covariance scalar. By varying the scalar from 0 (no variation) to 1 (natural-scene-typical variation), we examined how background variation affects performance in the task. Figure 3 shows examples of images used in our psychophysical task for different choices of the covariance scalar.

**Figure 3:**
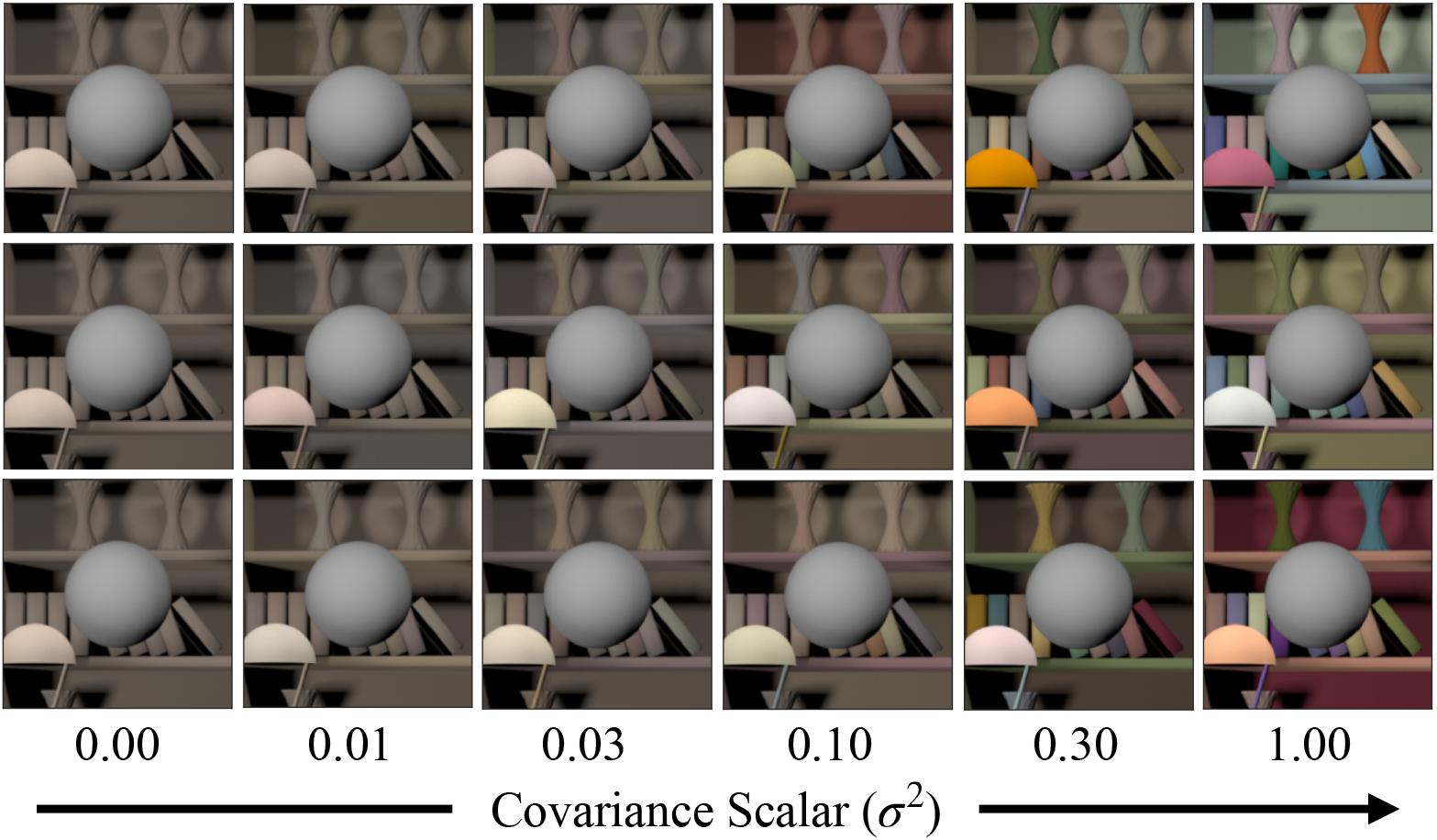
Variation in background object reflectances: The reflectance spectra of background objects were chosen from a multivariate normal distribution that modeled the statistics of natural reflectance spectra. The variation in the reflectance spectra was controlled by multiplying the covariance matrix of the distribution with a scalar. We generated images at six levels of the scalar. Each column shows three sample images at each of the six values of the scalar. The leftmost column corresponds to no variation and the rightmost column corresponds to the modeled variation of natural reflectances. The target object (sphere at the center of each panel) in each image has the same LRF. For each value of the scalar, we generated 1100 images, 100 each at 11 linearly spaced target LRF levels across the range [0.35, 0.45]. Discrimination thresholds were measured separately for each value of the covariance scalar.

The subsections below provide additional methodological detail.

### Preregistration

The experimental design and the method for extracting threshold from the data were preregistered before the start of the experiment. The preregistration documents are publicly available at: https://osf.io/7tgy8/.^4^

We preregistered three experiments. The first experiment (preregistered as Experiment 1) was abandoned because the task was too difficult. The findings of the second experiment (preregistered as Experiment 2 and referred to here as the control experiment) provide control data and are reported in the Appendix. In the body of the paper, we report preregistered Experiment 3 (referred to here as the main experiment). The details of the experimental methods below refer to preregistered Experiment 3; the methods for preregistered Experiment 2 were essentially the same with key differences (primarily the conditions studied) described in the Appendix.

A deviation from the preregistered plan for preregistered Experiment 2 was the change in the criteria to select observers for the experiment. The preregistered criterion for selecting an observer for this experiment was that an observer would be excluded if their mean threshold for the last two blocks in the practice session exceeded 0.025. After collecting data from 8 naive observers, we concluded that this criterion was too strict as only one observer met the criterion. Hence, we increased exclusion threshold from 0.025 to 0.030. The preregistered plans also indicated that each image would be presented for 500ms, but in the event we shortened this to 250ms.

We followed the procedure described in the preregistration document to extract threshold from the data. The document also indicated that the primary data feature of interest was the dependence of threshold on the covariance scalar and predicted that thresholds would increase with increasing background variability. The quantitative models of the data, however, were developed post-hoc.

### Reflectance and Illumination Spectra

The reflectance spectra for the background objects in the scene were generated using a model of naturally occurring surface reflectance spectra, as described in (Singh, Cottaris, Heasly, Brainard, & Burge, 2018). Briefly, we started with two datasets of surface reflectance functions (Kelly, Gibson, & Nickerson, 1943; Vrhel, Gershon, & Iwan, 1994) containing 632 surface reflectance measurements in total. The Kelly et al. dataset has 462 spectral measurements of Munsell papers, with each spectrum available to us (psychtoolbox.org) on wavelength support 400nm to 700nm at 5nm spacing. The Vrhel dataset has 170 spectral measurements, each spectrum measured in the wavelength range 390nm to 730nm at 2nm spacing. We converted to a common wavelength support of 10 nm spacing between 400nm and 700nm and combined the two datasets. We then used principal component analysis (PCA) to characterize the combined dataset. For this analysis, we mean centered the dataset and used the singular value decomposition (SVD) to obtain the eigenvectors of the mean-centered dataset. The reflectances in the mean-centered dataset were projected onto the eigenvectors to obtain their projection weights. The eigenvectors associated with the six largest eigenvalues captured more than 99.5% of the variance, so the rest of the analysis focuses on the projection weights on these eigenvectors. We approximated the empirical distribution of projection weights by a multivariate normal distribution. Reflectance spectra for the objects in the scene were generated by randomly sampling from this multivariate normal distribution and using the eigenvectors to construct samples of mean-centered surface reflectances. To these, we added back the mean of the surface reflectance dataset. We imposed a physical realizability constraint on the randomly generated spectral samples by ensuring that the reflectance at each wavelength was between 0 and 1. If the reflectance of a generated sample did not fall in this range at any wavelength, it was discarded.

The amount of variation in the surface reflectance of the background objects was controlled by multiplying the covariance matrix of the multivariate normal distribution (see above) by a covariance scalar. A covariance scalar of 0 corresponds to no background object reflectance variation. A covariance scalar of 1 corresponds to the full reflectance variation of the model of natural reflectance (Figure 3). We generated images for six logarithmically spaced values of covariance scalar: [0, 0.01, 0.03, 0.1, 0.3, 1.0]. Due to the physical realizability constraint, the actual variances of the projection weights for the generated spectral samples for some covariance scalars were lower than the corresponding variances of the underlying multivariate normal, and their distribution was not precisely multivariate normal.

The power spectrum of the light sources was chosen as that of standard daylight D65. We normalized the D65 spectrum by its mean power to obtain its relative spectral shape. This was multiplied by a fixed scalar with an arbitrarily chosen value of 5 to get the illuminant spectrum. This spectrum was used for all light sources in the visual scene and was not varied across the experiments reported here.

### Image Generation

The images were generated using software we refer to as Virtual World Color Constancy (VWCC) (github.com/BrainardLab/VirtualWorldColorConstancy). VWCC is written using MATLAB. It harnesses the Mitsuba renderer (Jakob, 2010) to render simulated images from scene descriptions, and also takes advantage of our RenderToolbox package (rendertoolbox.org; Heasly, Cottaris, Lichtman, Xiao, & Brainard, 2014). To render an image, we first create a 3D model that specifies the base scene. Objects and light sources can be inserted in the base scene at user specified locations. The 3D models utilized a base scene provided as part of RenderToolbox and modified using Blender, an open-source 3-D modeling and animation package (blender.org). Next, we assigned reflectance spectra and spectral power distribution functions to the objects and light sources in the scene (see Methods: Reflectance and Illumination Spectra). For each image, reflectances were assigned to the background objects by random draw from the reflectance model described above, with appropriate covariance scale factor. This procedure means that a set of images embodies the variation in background spectra described by the reflectance model, with each individual image containing a variety of background reflectances (Figure 3). Illumination spectra were not varied throughout the experiments reported here, and illumination spectra were as described in Methods: Reflectance and Illumination Spectra above.

Once the geometrical and spectral features were specified, we rendered a 2D multispectral image of the scene using Mitsuba, a physically-realistic open-source rendering system (mitsuba-renderer.org; Jakob, 2010). The images were rendered at 31 wavelengths equally spaced between 400nm and 700nm. The images were rendered with the camera field of view of 17° with an image resolution of 320-pixel by 240-pixels with the target object at the center. A 201-pixel by 201-pixel area, centered around the spherical target object, was cropped for display on the monitor.

To present the multispectral images on the monitor, they were first converted to LMS images using the Stockman-Sharpe 2° cone fundamentals (T_cones_ss2 in the Psychophysics Toolbox). Then the monitor calibration data and standard methods (Brainard, 1989; Brainard, Pelli, & Robson, 2002) were used to convert the LMS images to gamma corrected RGB images. A common scaling was applied to all images before rendering to ensure that they were within monitor gamut, so that the maximum linear channel RGB channel input was 0.9. The gamma corrected RGB images was presented on the monitor during the experiment.

### Stimulus Design

As noted above, we measured lightness discrimination thresholds for six values of the covariance scalar. For each value of the covariance scalar, we generated a dataset of 1100 images. The dataset had 100 images each at 11 values of the target object LRF. The LRF of the target object in the standard images was 0.4 and the LRF in the comparison image varied between 0.35 and 0.45 at steps of 0.01 (11 comparison levels). We generated 100 images at each comparison level, each with a different choice of the reflectance spectra of the background objects. The fact that we had 100 images for each target LRF allowed us to randomize the background object reflectances across the two intervals of each forced choice trial without excessive replication. For covariance scalar 0.00 we generated a set of 11 images, one at each LRF level, as the background remained fixed in this case. All images were generated without secondary reflections specified in the rendering process. The geometry of the 3D scene was also held fixed across all images.

When displayed on the experimental monitor, the average luminance of the standard image for covariance scalar 0.00 was 47.3 cd/m^2^. The average luminances of the target object for the 11 LRF levels were [67.0, 68.0, 68.9, 69.8, 70.7, 71.6, 72.5, 73.4, 74.2, 75.1, 75.9] cd/m^2^.

### Experimental Details

We define a trial as the presentation of two images (standard and comparison images) and collection of the observer’s response. We define an interval as the presentation of one of the images in the trial.

The experiment was structured as follows. We define a block of trials as the data collected at one covariance scalar with 30 trials at each of the 11 comparison levels. We define a permutation as a set of six blocks, where each block corresponds to one of the possible six covariance scalars. We collected three permutations for each observer, with a new random order drawn for each permutation. Thus, after the practice session (see Methods: Observer Recruitment and Exclusion), there were total 18 blocks. We divided these 18 blocks over 6 sessions, each session with 3 blocks. In each block, we randomly selected the images for the trials from the pre-generated image database. The first five trials of each block were moderate trials (as defined in Methods: Observer Recruitment and Exclusion) to acclimatize the observer to the experimental task. The responses for these five trials were not saved.

The trial sequence (comparison level, specific images, standard/comparison order) in a block was generated pseudo-randomly at the beginning of the block. For this, at each comparison lightness level, 30 standard and comparison images were chosen pseudo-randomly with replacement from the image dataset. The sequence of presentation of these 330 trials were randomized and saved. For each trial, the order of presentation of the standard and comparison image was also determined pseudo-randomly and saved. The trials were presented according to the saved sequence.

The trials in a block were presented in three sub-blocks of 110 trials each. At the end of each sub-block the observer took a break of minimum duration 1 minute. The observer could terminate the experiment anytime during the block. If an observer terminated a block, the data for that block was not saved. No observer terminated any block. One observer indicated a desire to postpone at the beginning of a session, due to fatigue for reasons unrelated to the experiment. The session was rescheduled.

At the beginning of the first experimental session (the practice session) for each observer, the experimenter explained the experimental procedures and obtained consent for the experiments. The experimenter then tested the observer for normal visual acuity and color vision. The observer was then taken to the experimental room, where the experimenter described the task, and the observer was shown the display, chin rest, and response box. The observer was dark adapted by sitting in the dark room for approximately 5 minutes. The observer then performed the familiarization block (see Methods: Observer Recruitment and Exclusion for explanation of familiarization block). After the familiarization block, the observer performed the other three blocks of the practice session. The practice session lasted about one hour.

Observers who met the inclusion criteria (see Methods: Observer Recruitment and Exclusion) then performed 18 blocks over 6 additional sessions, each on a separate day. The order of blocks for each observer was determined pseudo-randomly at the beginning of the practice session. As noted above, observers performed three blocks per session. Observers were dark adapted for 5 minutes at the beginning of each session. The data for all observers in the main experiment (preregistered Experiment 3) were collected over a period of four weeks.

Observers viewed the stimuli with both eyes.

### Observer Recruitment and Exclusion

Observers were recruited from the University of Pennsylvania and the local Philadelphia community and were compensated for their time. Observers were screened to have normal visual acuity (20/40 or better; with corrective eyewear, if applicable) and normal color vision, as assessed with pseudo-isochromatic plates (Ishihara, 1977). These exclusion criteria were specified in the preregistration document (see Methods: Preregistration). One observer was discontinued at this point for not meeting the normal visual acuity criterion.

Observers who passed the vision screening then participated in a practice session. This session also served to screen for observers’ ability to reliably perform the psychophysical task. At the beginning of the practice session, observers were familiarized with the task via a familiarization block. In the familiarization block, observers performed 40 trials of the task using images with covariance scalar 0.00 (10 easy trials, 10 moderate trials, and 20 regular trials). In the easy trials, the observers compared images with target object LRF 0.35 and 0.45. In the moderate trials, they compared images with target object LRF 0.40 to images with target object LRF 0.35 or 0.45. In the regular trials they compared images with target object LRF 0.40 to images with target object LRF in the range [0.35, 0.45]. The data from the familiarization block was not saved. The observer then performed three normal blocks for images with covariance scalar 0.00. At the end of the practice session, the mean threshold of the observer for the last two blocks was computed. The observer was excluded from further participation if their mean threshold for the last two blocks in the practice session exceeded 0.025 (log T^2^, −3.2). This exclusion criterion was specified in our preregistered protocol (See Methods: Preregistration).

Observers who met the performance criterion participated in the rest of the experiment.

### Observer Information

A total of 17 observers participated in the practice sessions for the control and main experiments (preregistered Experiments 2 and 3). To de-identify observer information in the data, observers were numbered in the order they performed the practice sessions. 10 observers participated in the practice sessions for the main experiment (preregistered Experiment 3; 6 Female, 4 Male; age 18-56; mean age 30.7). Four of these observers (Observer 2, Observer 4, Observer 8, and Observer 17) met the performance criterion set for screening (2 Female, 2 Male; age 23-56; mean age 38.25). All observers who advanced to the practice session had normal or corrected-to-normal vision (20/40 or better in both eyes, assessed using Snellen chart) and normal color vision (0 Ishihara plates read incorrectly). The visual acuities of the observers in the main experiment were: Observer 2, L = 20/30, R = 20/30; Observer 4, L = 20/15, R = 20/20; Observer 8, L = 20/30, R = 20/25; Observer 17, L = 20/20, R = 20/20. Observers 2, 8, and 17 wore personal corrective eyewear both during vision testing and during the experiments. Observer 4 did not require or use corrective eyewear.

### Apparatus

The stimuli were presented on a calibrated LCD color monitor (27-in. NEC MultiSync PA271W; NEC Display Solutions) in an otherwise dark room. The monitor was driven at a pixel resolution of 1920 x 1080, a refresh rate of 60Hz, and with 8-bit resolution for each RGB channel. The host computer was an Apple Macintosh with an Intel Core i7 processor. The experimental programs were written in MATLAB (MathWorks; Natick, MA) and relied on routines from the Psychophysics Toolbox (http://psychtoolbox.org) and mgl (http://justingardner.net/doku.php/mgl/overview). Responses were collected using a Logitech F310 gamepad controller.

The observer’s head position was stabilized using a chin cup and forehead rest (Headspot, UHCOTech, Houston, TX). The observer’s eyes were centered horizontally and vertically with respect to the display. The distance from observer’s eyes to the monitor was 75cm.

### Monitor Calibration

The monitor was calibrated using a spectroradiometer (PhotoResearch PR650). To calibrate the monitor, we focused the spectroradiometer on a patch displayed on the center of the monitor. The patch size was 4.66cm x 4.66cm (3.56° x 3.56°). The optics of the radiometer sampled the emitted light from a 1° circular spot within the patch. The spectral power distribution of the three monitor primaries was measured in the range 380nm to 780nm at 4nm steps. The gamma functions for each primary were determined from measurements of the spectral power distribution for each primary at 26 equally spaced input values for that primary, in the range [0, 1] where 1 corresponds to the maximum input value of the device. These gamma functions as well as the light emitted by the monitor for an input of 0 were accounted for in the stimulus display procedures. The spectral power distribution was also measured for 32 different combinations of RGB input values. These measurements were used to check the performance of the display. The maximum absolute deviation of the x-y chromaticity between the measured values and those predicted from the calibration was 0.0028 and 0.0027 for x and y chromaticity respectively, and less than 1% for luminance.

### Stimulus Presentation

The size of each image was 2.6cm x 2.6cm on the monitor, corresponding to 2° by 2° visual angle. The target object size on the screen in the 2D images was ~1° in diameter. Each image was presented for 250ms (this was a deviation from the preregistration document, which specifies the presentation time as 500ms), with an inter-stimulus interval of 250ms and inter-trial interval of 250ms. Inter-stimulus interval (ISI) is defined as the interval between the first and the second image presented on each trial. The response for each trial was collected after both the images had been displayed and removed from the screen. The observer could take as long as they wished before entering the response. Feedback was provided via tones presented after the response to allow observers to maximize their performance. The next trial was presented 250ms (ITI) after the feedback. Thus, the actual inter-trial interval depended on the response time of the observer.

### Psychometric Function

The proportion comparison chosen data was used to obtain the psychometric function for each block. Each block consisted of 330 trials with 30 trials at each comparison lightness level. At each lightness level, we recorded the number of times the observers chose the comparison image to be lighter. The proportion comparison chosen data were fit with a cumulative normal using the Palamedes toolbox (Prins & Kingdom, 2018) to obtain four parameters of the psychometric function: threshold, slope, lapse rate and guess rate. The lapse rate was constrained to be equal to the guess rate and to be in the range [0, 0.05]. The psychometric function was fit using the maximum likelihood method. The threshold was obtained as the difference between the LRFs at proportion comparison chosen 0.76 and 0.50 as obtained from the cumulative normal fit.

### Ethics Statement

All experimental procedures were approved by University of Pennsylvania Institutional Review Board and were in accordance with the World Medical Association Declaration of Helsinki.

### Code and Data Availability

For each observer, the proportion comparison chosen data for the 18 experimental blocks as well as the thresholds are provided as supplementary information (SI). The SI also provides the MATLAB scripts to generate Figures 2, 4, 5, 6 and 7 and the scripts to obtain thresholds of the linear receptive field formulation of the model (model described below). The computed retinal images used as input to the model are provided as .mat files in a zip folder. The SI is available at: https://github.com/vijaysoophie/EquivalentNoisePaper.

**Figure 4:**
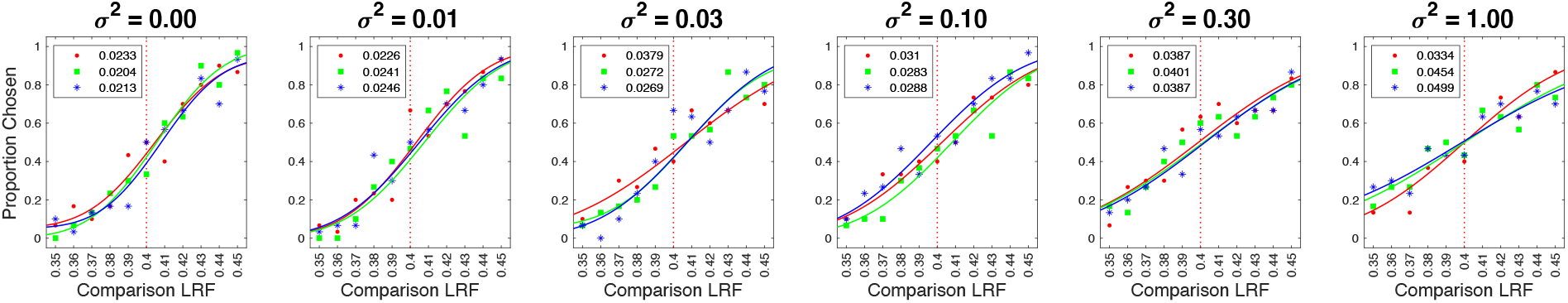
Psychometric functions for Observer 2. We measured the proportion comparison chosen data at six values of the covariance scalar (*σ*^2^), separately in three blocks for each observer. The data for each block was fit with a cumulative normal to obtain the discrimination threshold (see Figure 2). Each panel plots the measured values and the cumulative fit to the proportion comparison data for each of the three blocks, for Observer 2. The values in the legend provide the estimate of lightness discrimination threshold for each block obtained from the cumulative fit. See Figure S3 for the psychometric functions of all observers.

**Figure 5:**
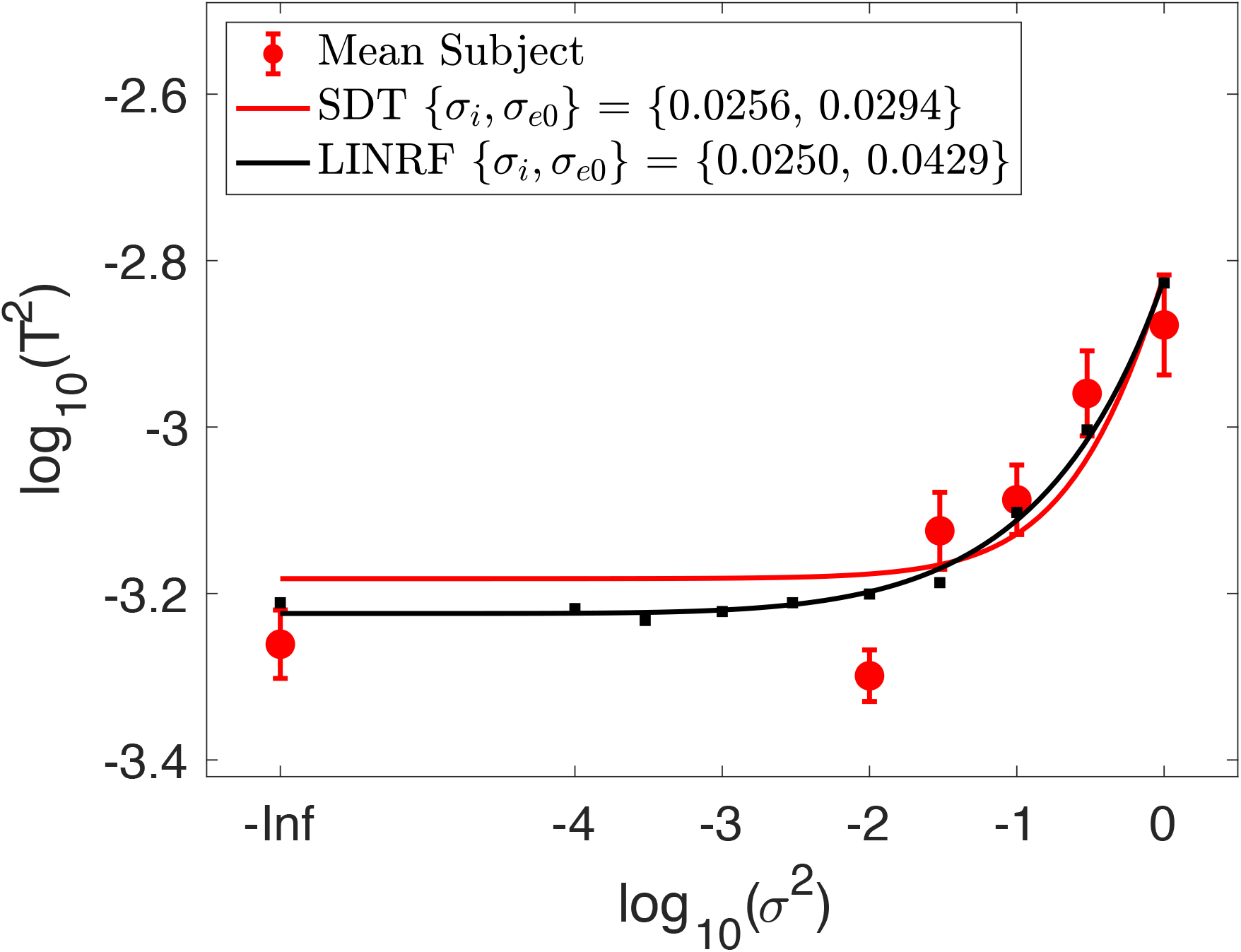
Background variation increases lightness discrimination threshold. Mean (N = 4) log squared threshold vs log covariance scalar from the human psychophysics (red circles). The error bars represent +/- 1 SEM taken between observers. The fit of the STD formulation of the model (Equation 4) is shown as the red curve. The parameters corresponding to this fit are provided in the legend. The threshold of the fit linear receptive field (LINRF) formulation was estimated by simulation at 10 logarithmically spaced values of the covariance scalar (black squares). The black smooth curve is a smooth fit to these points of the functional form log_10_ *T^2^* = *a* + *b*^(*x*+*c*)*d*^ where *x* = log_10_ *σ*^2^ and *a, b, c* and *d* are parameters adjusted in the fit. This functional form was chosen simply to provide a smooth curve through the simulated thresholds and has no theoretical significance. The parameters of the LINRF fit are also provided in the legend.

**Figure 6:**
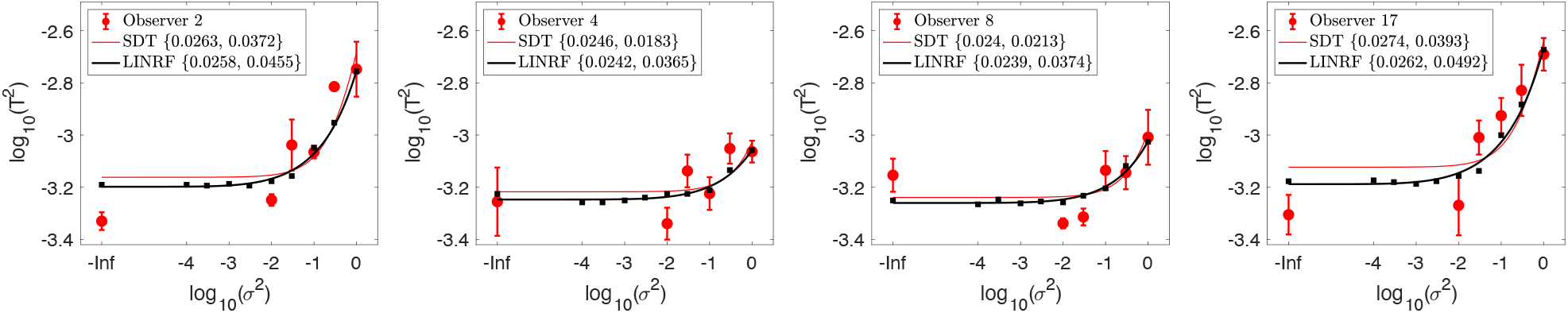
Threshold of individual human observers. Mean (across sessions) squared threshold vs log covariance scalar for individual human observers. Same format as Figure 5; here the error bars represent +/- 1 SEM taken across the three blocks for each observer. The parameters of the SDT and LINRF formulations were obtained separately for each observer and are provided in the legend, in order 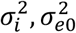.

**Figure 7:**
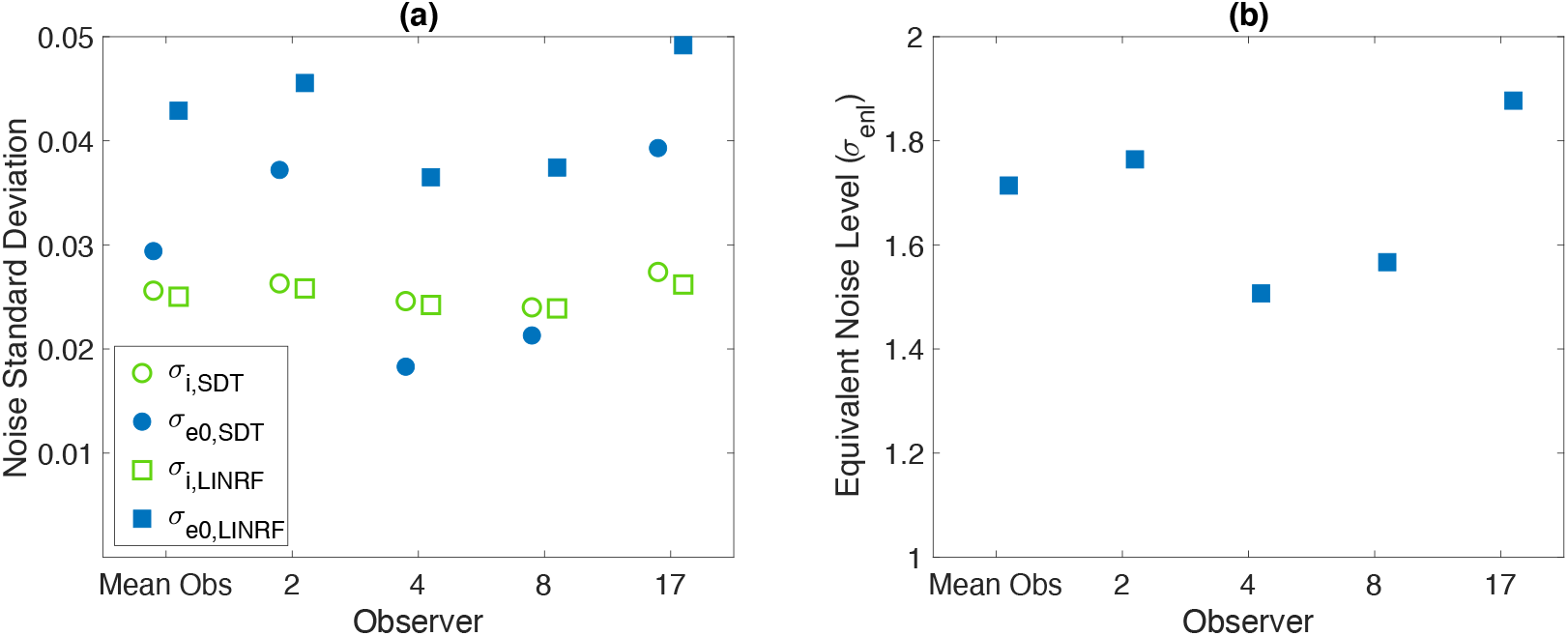
Equivalent noise analysis. **(a)** The left panel shows the parameter estimates for the two model formulations for the mean data and each individual observer. From these, we can estimate the equivalent noise level (*σ_enl_*) for background object reflectance variation corresponding to the full model of natural reflectance variation (covariance scalar *σ*^2^ = 1). **(b)** The equivalent noise level is provided for the mean data and each individual observer in the right panel.

## 3 MODEL

The data collected in the experiments characterize how lightness discrimination thresholds increase with the variance of a task-irrelevant stimulus variable. Interpreting the data is aided by a model that relates the changes in discrimination thresholds to the underlying precision of the perceptual representation. The model provides a way to connect the variance of a task-irrelevant property to the precision of the perceptual representation of the task-relevant stimulus variable (here lightness). The model we employ shares features of models that have been used to understand how contrast thresholds are elevated in the presence of contrast noise (see e.g., Legge, Kersten, & Burgess, 1987; Pelli, 1990). We provide a full development of the model here, however, as the current application of the underlying ideas differs substantially from previous applications.

We first introduce an analytic formulation, derived in the context of signal detection theory (SDT formulation). We then show how this can be instantiated as a linear receptive field model whose performance can be simulated (LINRF formulation). An important advantage of the LINRF formulation is that it can accommodate the physical-realizability constraint incorporated into our statistical model of naturally occurring reflectances.

The model allows us to express the variation of the task-irrelevant stimulus variable in units of equivalent noise standard deviation, where an equivalent noise standard deviation of 1.0 corresponds to the amount of external variation whose effect on the perceptual representation of the task-relevant stimulus variable is the same as that of the intrinsic internal noise that limits discrimination in the absence of task-irrelevant external variation. In this way, we can understand the effect of the task-irrelevant variability on thresholds in perceptually meaningful units of equivalent noise level. Task-irrelevant variability with an equivalent noise level less than one have little impact on the visual system, since its effects are dominated by intrinsic variability. Levels of task-irrelevant variability with an equivalent noise level greater than one do intrude on perception. The equivalent noise level indicates the magnitude of the intrusion in units that connect to intrinsic precision. Equivalent noise is similarly used in the literature on contrast noise masking (again see e.g., Legge, Kersten, & Burgess, 1987; Pelli, 1990).

### SDT Model Formulation

We first formulate our model in the context of signal detection theory (Green, 1966). We model the visual response to the target object in each image by a univariate internal representation denoted by the variable *z*. This variable depends on the image and is perturbed by noise. We assume that for any fixed image, *z* is a normally-distributed random variable whose mean depends on the target object LRF. For each image, we assume that *z* is perturbed on a trial-by-trial basis by independent zero-mean normally-distributed noise, and we assume that the variance of this noise is the same for the response to all images. We refer to the noise that perturbs *z* for a fixed image as the internal noise and denote its variance as 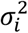. For each trial of the experiment, *z* takes on two values, *z_s_* and *z_c_*, one for the interval containing the standard and the other for the interval containing the comparison.

If we consider performance for a particular pair of target standard and comparison LRFs, performance depends both on the difference between the expected values of *z* for each pair of LRFs, *μ_s_* and *μ_c_*, and on the value of 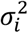. In our experimental design we have ensembles of images with different backgrounds, for each value of the target object LRF and background covariance scalar. The fact that we draw stochastically from these ensembles on each trial introduces additional variability into the value of the decision variable *z* that corresponds to a fixed target LRF. We call this the external variability, and model it as a normal random variable with zero mean and variance 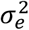. We assume that 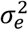 depends on the experimentally chosen covariance scalar, but not on the target sphere LRF. Thus, the distributions of *z_s_* and *z_c_*, for a particular choice of target standard and comparison LRF and covariance scalar, are given by *P*(*z_s_*) = *N*(*μ_s_, σ_t_*) and *P*(*z_c_*) = *N*(*μ_c_, σ_t_*). Here *μ_s_* is the mean value of the internal representation to the standard image and *μ_c_* is the mean value of the internal representation to the comparison image. The overall standard deviation *σ_t_* is obtained via 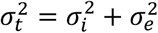, where 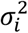 and 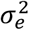 are the variance of the internal and external noise.

For a 2AFC discrimination task in the context of signal detection theory, the observer makes their decision based on a comparison of *z_s_* and *z_c_*, choosing the interval with the higher value of *z* as that with the higher stimulus value. The observer’s sensitivity depends on the mean values and the variance of *z*, and is captured by the quantity d-prime: *d*′ = (*μ_c_* – *μ_s_*)/*σ_t_*. D-prime measures the distance between the two distributions in standard deviation units. A value of *d*′ = 0 corresponds to an inability to distinguish between the standard and the comparison image. Larger values of *d*′ indicate increasing discriminability.

For a fixed value of *d*′, the difference in mean values is directly proportional to the standard deviation *σ_t_*:

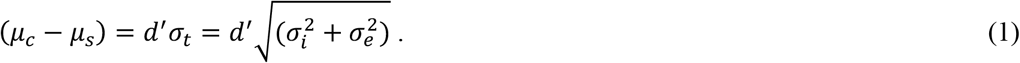

We further assume that the difference in mean value of the internal variable (*μ_c_* – *μ_s_*) is proportional to the difference in the LRFs of the target object in the standard and comparison images (Δ_LRF_). That is, (*μ_C_* – *μ_s_*) *C* Δ_LRF_, where *C* is the proportionality constant. This yields

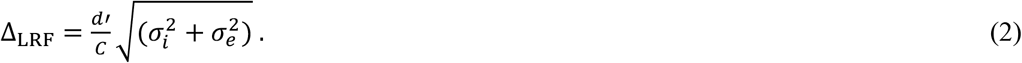

When we measure threshold in a 2AFC task, we choose a criterion proportional correct and find the Δ_LRF_ that corresponds to that proportion correct. Our choice of 0.76 corresponds to *d*′ = 1. In addition we can choose *C* = 1, in essence setting the units for *z* to match those of the target LRF.

In our experiment, external variability was induced by changing the reflectance of the objects in the background. We used a multivariate normal distribution to generate the reflectance functions of the background objects.^5^ To change the amount of external noise, we scaled the covariance of the multivariate normal distribution by multiplying its covariance matrix with a scalar. Thus, for our experiments we have

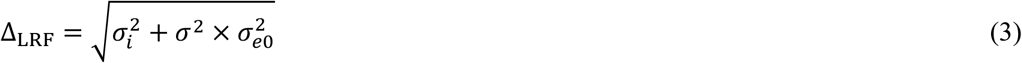

where *σ*^2^ is the covariance scalar and 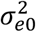 is the external noise introduced when the ensemble of images for each value of target LRF has the reflectance of the background objects drawn from our model of natural reflectances.

Converting the equation above to the form we use to represent the data, we have

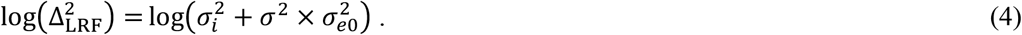

The equation above predicts that the form of threshold 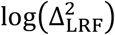 as a function of covariance scalar *σ*^2^ should increase monotonically. For small values of 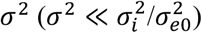, the threshold will approach a constant giving 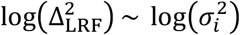. For large values of 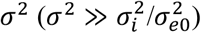, the quantity 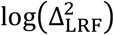 will approach a straight line with slope 1 in the 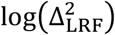 versus log(*σ*^2^) plot. Fitting the measurements with Equation 4 allows us to check whether the model describes the data, as well as to determine the two parameters 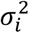 and 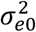. In particular, we can establish the relative contribution of the internal representational variability and external stimulus variability in limiting lightness discrimination. The parameter 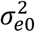 quantifies how much the variation in background object reflectances intrudes on the internal representation *z* that mediates the lightness discrimination task. The value of 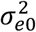 may be compared directly to the intrinsic precision of that representation characterized by 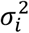.

### Equivalent Noise Level

The SDT formulation allows us to introduce the concepts of equivalent noise and equivalent noise level. The equivalent noise is the amount of external variation that has the same effect on the decision variable *z* as the internal noise. The external variation is characterized experimentally by the covariance scalar (together with the underlying model of natural reflectances which is held fixed across the experiments). Once the model parameters 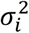 and 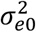 are determined from the data, we can find the covariance scalar 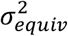 that produces externally-generated equivalent noise

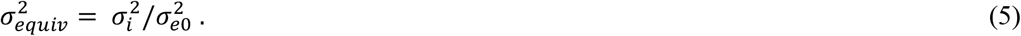

This in turn allows us to express the covariance scalars in terms of their equivalent noise level, which gives their effect on the perceptual representation relative to the effect of the internal noise. Thus

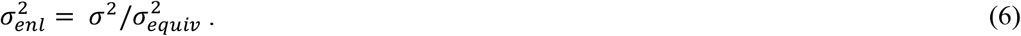

For 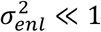, the effect of the external noise is negligible and does not affect the perceptual representation and the internal noise dominates the precision of the representation. For 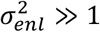, the effect of the external noise dominates the perceptual representation, and the visual system has not insulated the representation of the task-relevant stimulus variable from the variation in the task-irrelevant perceptual variable. When the equivalent noise level is ~1, the effect of the external variability is matched to that of the internal variability. At this operating point, further insulation of the task-relevant representation will not lead to significant further increases in the precision of this representation. We can thus use the equivalent noise level as a calibrated metric for assessing the magnitude of the perceptual effect of various levels of task-irrelevant stimulus variation.

### Linear Receptive Field Formulation

When external noise added to the images is characterized by a multivariate normal and the decision noise is normal, a simple linear receptive field (LINRF) formulation is equivalent to the SDT formulation developed above. We develop this equivalence below. The advantage of the LINRF formulation is that it can easily be applied directly to images and to cases where the internal or external variability is non-normal. In our application, there are two non normalities. First, although the projection weights for linear model of naturally-occurring reflectance are drawn from a multivariate normal distribution, the constraint that the resulting reflectance functions lie within the range between 0 and 1, implemented to satisfy physical realizability, makes the overall distribution non-normal. Second, we incorporate into the model the Poisson variability of the cone excitations.

We begin with development that connects the LINRF formulation to the SDT formulation. In the LINRF formulation, the decision variable is computed from the displayed stimulus as the response of a single unit whose responses are a linear function of the stimulus image. Denote the stimulus image by the column vector *I*, and the receptive field by the column vector *R*. The entries of *I* are the radiant power emitted by the monitor at each image location. The entries of *R* are the corresponding sensitivities of the linear receptive field to each entry of *I*. The response of the receptive field is given as *r_i_* = *R^T^l* + *η_i_*, where *η_i_* is a random variable representing a draw of zero mean normally-distributed internal noise (variance 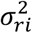) in the receptive field response for a fixed image. We assume that 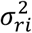 is independent of *I*.

Denote *I*_*s*0_ and *I*_*c*0_ as the standard and comparison images without external noise. External normally-distributed noise is added to both *I*_*s*0_ and *I*_*c*0_, with covariance matrix Σ_*e*_. The external noise need not have zero mean. After incorporation of the external noise, the response of the receptive field to the comparison and standard images is given by

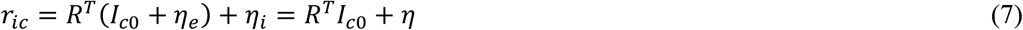

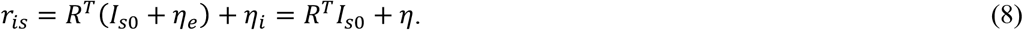

Here *η_e_* is a random variable representing a draw of external noise, *η_i_* represents the internal noise, and *η* is a random variable representing the overall effect of the external and internal noise. Since the receptive field and noise models are linear and normal, *η* is normal with variance

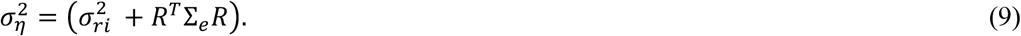

The mean difference between the receptive field response to the comparison and the standard image is given by (*μ_c_* – *μ_s_*) = *R^T^* (*I*_*c*0_ – *I*_*s*0_) = *C*′Δ_LRF_. Here *I*_*s*0_ and *I*_*c*0_ are the standard and comparison images without external noise added, *C*′ is a constant, and Δ_LRF_ is as defined is the SDT section above. The second equality follows because 1) the difference between *I*_*c*0_ and *I*_*s*0_ is proportional to Δ_LRF_ as only the target LRF changes between these two images and 2) even if the mean of the external noise is non-zero, its effect cancels when we obtain the mean difference in response.

We associate the linear receptive field response with the internal representation *z* of the SDT formulation developed above. That is, we assume that on each trial, the observer chooses as lighter the interval for which the response of the receptive field is greater. Following the development of the SDT formulation, we have

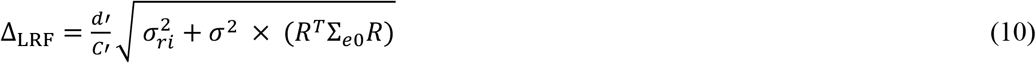

where we have introduced the covariance scalar *σ*^2^ in the term corresponding to the variance of the external noise, and where Σ_*e*0_ denotes the covariance matrix of the external noise corresponding to the level of variation in natural images. Comparing to relation derived in the SDT model (Equation 3), we see that this is the same functional form for the relation between Δ_LRF_ and *σ*^2^ as derived there, where we associate 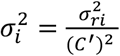 and 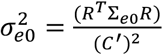.

To fit the LINRF formulation and relax its assumptions, we compute how images produce retinal cone excitations and employ a one-parameter description of a simple center-surround receptive field that draws upon the output of the cones. We use simulation to compute model responses for any choice of 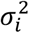. This procedure is described in more detail below. Once the fitting procedure establishes *R* and 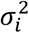 that best account for the data, we then find 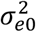 directly by passing the images corresponding to *σ*^2^ = 1 through the receptive field and finding the resulting variance. These parameters in turn allow us to compute 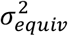 and 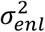 for the LINRF formulation.

### Fitting the SDT Model Formulation

The model was fit to the threshold versus covariance scalar data to obtain the parameters 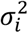 and 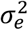. The parameters were obtained by minimizing the mean squared error between the measured and predicted threshold using the MATLAB function *fmincon.* The best fitting parameters were estimated separately for the mean observer and the individual observers.

### Fitting the Linear Receptive Field Model Formulation

We fit the linear receptive field (LINRF) model using a simulation approach. We used simulation for two reasons. First, it allows us to incorporate a model of the early visual system into the computations.

Second, it provides a way to account for truncation in the normally-distributed model of natural reflectances.

The model of initial visual encoding was as described by Singh et al. (2018), and was implemented using the software infrastructure provided by ISETBio (ISETBio; isetbio.org; Cottaris, Jiang, Ding, Wandell, & Brainard, 2019). It incorporated typical optical blur (Thibos, Hong, Bradley, & Cheng, 2002) and the Poisson noise that perturbs cone photoreceptor isomerizations in the retina (Rodieck, 1998). In addition, it included axial chromatic aberration (Marimont & Wandell, 1994), and spatial sampling by the mosaic of long (L), middle (M) and short (S) wavelength-sensitive cones (Brainard, 2015). The L:M:S cone ratio in the cone mosaic was chosen to be 0.6:0.3:0.1 (1523 L-cones, 801 M-cones, 277 S-cones). The CIE physiological standard (CIE, 2007), as implemented in ISETBio, was used to obtain LMS cone fundamentals. Cone excitations were calculated as the number of photopigment isomerizations in a 100ms integration time, and included simulation of the Poisson variability of the isomerizations (Rodieck, 1998). The cone isomerizations were demosaiced using linear interpolation to estimate LMS isomerization images. Further, the isomerizations of each cone class was normalized by the summed (over wavelength) quantal efficiency of the corresponding cone class, to make the magnitude of the signals from the three cone classes similar to each other. This normalization occurred after incorporation of Poisson noise and did not affect the signal-to-noise ratio of the signals from the different cone classes.

The dot product of the LMS isomerization images was taken with a simple center-surround linear receptive field. The receptive field was square in shape to match the image size. Its center was a circle of radius equal to the size and at the location of the target object in the image. The central region was taken to have spatially-uniform positive sensitivity, while the surround was taken to have spatially-uniform negative sensitivity. Each point in the central region had sensitivity v_c_ = 1, and each region of the surround had sensitivity denoted by v_s_. The RF was the same for each of the three cone classes. The RF response was taken as the sum of the L, M and S RF component responses. Normally-distributed internal noise with zero mean was added to the resulting dot product. The variance of the internal noise (*σ_ri_*) and the value of the RF surround sensitivity (v_s_) were the two parameters of the model.

The threshold predictions of the LINRF formulation for any choice of model parameters were obtained using simulation of a two-interval force choice paradigm similar to the experiment. For each trial, we randomly sampled a standard image and a comparison image from our dataset, following the procedure used in the experiment. We obtained the response of the receptive field (noise-added dot product) to the images and compared them to determine the simulated choice on that trial. This process was repeated 10,000 times for each of the 11 comparison LRF levels. The proportion comparison chosen data were used to fit the psychometric function and obtain the discrimination threshold, similar to the method used for the human psychophysical data. We estimated model threshold for the six values of covariance scalar at which we performed the human experiments.

We calculated the mean squared error (averaged over the six covariance scalar values) between the thresholds of the human data being fit and the computational model for a large set of values of the two model parameters: the variance of the decision noise (*σ_ri_*) and the value of the RF surround (v_s_). The mean squared error values obtained as a function of these two parameters were fit with a degree two polynomial of two variables using the MATLAB *fit* function. The resulting polynomial was evaluated to estimate the parameters with lowest mean square error. These parameters were then used to estimate the internal and external noise standard deviation of the LINRF formulation using the relations: 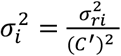 and 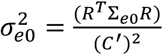 as explained above, where the constant *C*′ was obtained by solving *R^T^*(*I*_*c*0_ – *I*_*s*0_) = *C*′Δ_LRF_.

The best fitting parameters were estimated separately for the mean observer and the individual observers.

## 4. RESULTS

### Human Lightness Discrimination Thresholds Increase with Background Reflectance Variation

We measured lightness discrimination thresholds of human observers as a function of the amount of variation in the reflectance spectra of the background objects in the scene. The amount of variation was determined by the covariance matrix of the multivariate normal distribution from which the spectra were sampled. We controlled the variance by multiplying the covariance matrix by a covariance scalar (*σ*^2^). We measured discrimination thresholds of four human observers at six values of the covariance scalar. The threshold was measured three times (three separate blocks) for each observer and for each value of covariance scalar. The psychometric functions for each block/covariance scalar value are shown for one observer in Figure 4 and for all observers in Figure S3. Inspection of the psychometric functions shows that their slopes steadily decrease with increasing covariance scalar, corresponding to an increase in thresholds.

Figures 5 and 6 show the data in more digested form. These plots show explicitly how the discrimination thresholds change with the amount of variability in the reflectance of the background objects. In Figure 5, mean log threshold squared (averaged across observers, N = 4) is plotted against the log of the covariance scalar. Figure 6 plots thresholds in the same format for the individual observers, with the data averaged over the three blocks for each covariance scalar. The choice to plot the data as log threshold-squared against the log of the covariance scalar was motivated by the relatively simple expression of the SDT model formulation’s predictions for this representation (see Equation 4 and following text). Table S2 provides the thresholds and SEMs from Figure 6 in tabular form.

For low values of the covariance scalar, the thresholds are nearly constant and are similar across observers. As the covariance scalar increases, log squared threshold rises. These features are seen in the mean data (Figure 5) and in the data for all observers (Figure 6). The covariance scalar value at which thresholds start to increase is also similar across observers. There is some individual variability, however, in the slope of the rising limb of the measured functions.

### Modeling the Impact of Background Reflectance Variation

To interpret the data further, we fit the data with two formulations of our model (see Model section above). The performance of both the SDT and LINRF model formulations is determined by two fundamental factors. The first factor is variability in the perceptual representation of lightness internal to the visual system (i.e., internal noise, model parameter 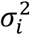). The second factor is the effect of experimentally-induced task-irrelevant stimulus variability (i.e., background object reflectance variability) on the same perceptual representation (i.e. external noise, model parameter 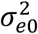). Roughly speaking, threshold with no external variation (covariance scalar *σ*^2^ = 0) establishes the level of the internal noise, while the way threshold increases with covariance scalar determines 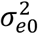. The fits determine the parameters of the model as well as allow us to examine how well the model fits the data.

The fits to the mean observer data are shown in Figure 5; the fits to the individual observer data are shown in Figure 6.

The fit of the analytic STD formulation (red curves) captures the main trends in the mean data and similarly for the fits to the individual observer data. Detailed examination, however, reveals that this formulation tends to overestimate thresholds in the low covariance scalar regime. An alternative way of putting this is that it underestimates the rising slopes as covariance scalar increases. Because the rising slope of this formulation asymptotes to 1, the SDT formulation of the model is not able to simultaneously describe thresholds over the full covariance scalar range.

The fits of the LINRF formulation (black curves) are better. The fit to the mean data does an excellent job of capturing these data, and the fits to the individual observer data are also improved relative to the SDT formulation. We attribute the improvement in fit of the LINRF formulation primarily to its ability to account for the truncation of our experimental reflectance distributions, which the SDT formulation cannot do (see Section 3 Model).

The model fits provide estimates of internal and external noise for the human observers in this task. Figure 7 (left panel) plots the estimates of the internal and external noise standard deviations (quantities *σ_i_* and *σ*_*e*0_, for both the SDT model and the LINRF formulation. There is good consistency in the value of *σ_i_* across observers, the model’s manifestation of the observations that thresholds for low covariance scalars are similar across observers. There is more variability in *σ*_*e*0_ across observers, corresponding to the individual variability seen in the rising limb of the threshold versus covariance scalar plots.

The Poisson noise included in the LINRF formulation does not typically limit human discrimination performance at daylight light levels (Banks, Geisler, & Bennett, 1987; Cottaris, Jiang, Ding, Wandell, & Brainard, 2019). Thus, it is not surprising that the mean values of the internal noise standard deviation parameter σ¿ for the LINRF formulation are close to those obtained with the SDT formulation (SDT formulation: mean value of internal noise standard deviation across observers 0.0256, value from fit to mean data 0.0256; LINRF formulation: mean value of internal noise standard deviation across individual observers 0.0250, value from fit to mean data, 0.0250).

The estimates of the external noise standard deviation parameter *σ*_*e*0_ are higher for the LINRF formulation than for the SDT formulation (SDT formulation: mean value of external noise standard deviation 0.0290, value from fit to mean data across observers 0.0294; LINRF formulation: mean value of external noise standard deviation across observers 0.0421, value from fit to mean data, 0.0429). This is consistent with the observation that the SDT formulation underestimates the rise in thresholds with increasing covariance scalar, while this rise is captured more accurately by the LINRF formulation, presumably because the latter incorporates the constraint that the reflectance values at each wavelength are physically realizable (i.e., reflectances lie between 0 and 1).

If we focus on the estimates from the better fitting LINRF formulation, we can compute the equivalent noise level (*σ_enl_*) corresponding to covariance scalar *σ*^2^ = 1, the level of background object reflectance variation corresponding to our full model of natural reflectance. For the fits to the mean data, this equivalent noise level is ~1.7. This as well as values for the individual observers are plotted in the right panel of Figure 7. This tells us that, for our experimental conditions, the variability in the human representations of lightness induced by naturally occurring variation in background object reflectances is within a factor of two of the limits imposed by the intrinsic precision of that representation. Had the value been closer to 1, we would have concluded that the visual system had discounted the effect of variation in the background object reflectances about as required, given the intrinsic precision of the lightness representation. The fact the equivalent noise level is higher than 1 but not tremendously so is consistent with the idea that the visual system has a degree of lightness constancy, but that this constancy can be incomplete (see e.g., Gilchrist, 2006; Kingdom, 2011; Murray, 2021).

## 5 DISCUSSION

The perceived lightness of an object can depend on the scene in which it lies. Stabilization of the lightness representation against variation in scene properties extrinsic to the object’s surface reflectance is referred to as lightness constancy. In this paper, we introduced a new psychophysical approach for characterizing lightness constancy. The approach is based on measuring how lightness discrimination thresholds vary with experimentally introduced variation in scene properties extrinsic to the object’s reflectance. Specifically, we studied how lightness discrimination thresholds are impacted by variation in the reflectance of the background objects in naturalistic scenes rendered using computer graphics. Our results (Figures 5 and 6) show that when the variation in the reflectance of background objects is small, discrimination thresholds are nearly constant. In this regime, performance is limited primarily by internal noise. As the amount of background object reflectance variation increases, the effect of external variation starts dominating that of the internal noise, and discrimination thresholds increase. We analyzed the data using a modeling approach used previously to study effect of external noise on contrast detection (Legge, Kersten, & Burgess, 1987; Pelli, 1990; Pelli & Farell, 1999). This approach allows us to relate the effect of background object reflectance variation to the intrinsic precision of the lightness representation. The intrinsic precision depends on the observer’s internal noise, which limits performance in the absence of external variation. The model compares discrimination thresholds with and without extrinsic variations to quantify variance in the perceptual representation of lightness induced by extrinsic variation. It allows us to express the effect of extrinsic variation as an equivalent noise level (*σ_enl_*), that is relative to the standard deviation of the intrinsic noise. In this way, we use the intrinsic noise as a benchmark to interpret the magnitude of the equivalent noise from the external variation. We find that the effect of the external variability introduced by variation of background object reflectances in naturalistic scenes is within a factor of two of the intrinsic precision of the lightness representation. More generally, our work provides a method to quantify the effect of variation in a task-irrelevant properties on the perception of task-relevant property, and is thus applicable to understanding other perceptual constancies beyond the lightness constancy we focused on here.

### Relation to Contrast Detection in Contrast Noise

As noted, our paradigm and model have conceptual roots in the literature on contrast detection in contrast noise. The concept of equivalent noise plays an important role in this literature (Legge, Kersten, & Burgess, 1987; Pelli, 1990; Pelli & Farell, 1999). However, there is an important difference between the way the ideas are applied to understand contrast detection and the way we have leveraged them here. In the contrast detection literature, detection in the absence of external noise is conceptualized as limited by two distinct factors. One factor is the internal variability in the observer’s representation of contrast. The other factor is the efficiency with which the observer’s decision processes makes use of the information provided by this representation, which is inferred through an ideal observer analysis applied to high external noise conditions, where effects of internal noise are swamped by those of the external noise (Pelli, 1990; Pelli & Farell, 1999). This separation is enabled when such an ideal observer calculation is available, and in practice is more straightforward when the stimulus being detected/discriminated and the external noise being added have commensurate units (e.g. contrast energy). In our work, the task-relevant and task-irrelevant stimulus variables vary along distinct dimensions of the stimulus space (e.g., affect distinct image locations). Currently we do not have in hand an ideal observer calculation that would allow us to compute the visual system’s efficiency in using the available information. Obtaining and integrating such a calculation would be of interest. Singh, Cottaris, Heasly, Brainard, & Burge (2018) provide a possible approach, but employing that approach would require measurements with a larger set of task-irrelevant variation (e.g., illumination as well as background) than available from the current data.

### Spatial and Chromatic Properties of the Stimuli

We used small image patches in our study. The small size of the image patches is a notable difference between our stimuli and natural viewing. In this initial deployment of our paradigm, we thus focused on effects of background object reflectance variation that are nearby the test object. The observed effects may be mediated by relatively small populations of neurons. The use of small image patches is not a necessary requirement of our paradigm, which could be extended to larger images. Such extension could reveal additional effects not captured by the current experiments.

In addition to using small patches, we did not vary the spatial structure of the array of objects in the rendered scenes. Manipulating spatial structure, in addition to increasing image size, may provide a way to use our paradigm to measure the spatial tuning of the mechanism(s) mediating the background effect. This approach is loosely analogous to how manipulating the structure of contrast noise may be used to examine the tuning of mechanisms supporting the detection of contrast-defined targets (Henning, Hertz, & Hinton, 1981; Rovamo, Franssila, & Nasanen, 1992; Losada & Mullen, 1995; Nachmias, 1999; Rovamo, Raninen, & Donner, 1999).

Although we restricted our measurements to lightness discrimination thresholds, our variation of the reflectance properties of the background objects was not limited to variation in overall reflectance. The choice to introduce background object reflectance variation along more spectral dimensions (affecting e.g. background object hue and saturation) than used for target object variation was somewhat arbitrary – we could have restricted the background object reflectance variation to one dimension (e.g. overall scale of reflectance spectra) or studied discrimination of additional (e.g. chromatic) dimensions of target object variation. As with the case of spatial structure above, extending the measurements to a wider range of stimuli is of interest. Indeed, it may be possible to manipulate the chromatic structure of the variation in background object reflectances with the goal of understanding the chromatic tuning of the background object reflectance variation’s effect on the lightness discrimination thresholds, as well as on other target object discriminations. This would again be analogous to how noise-based approaches have been used to characterize chromatic tuning of mechanisms that support the detection of chromatically-defined contrast targets (Gegenfurtner & Kiper, 1992; Sankeralli & Mullen, 1997; Giulianini & Eskew, 1998; Monaci, Menegaz, Süsstrunk, & Knoblauch, 2004).

### Link Between Thresholds and Suprathreshold Perceptual Judgments

The technique developed here probes the constancy of a perceptual representation of a task-relevant variable (e.g., perceived object lightness) by measuring how variation in a task-irrelevant scene variable (e.g., background object reflectances) elevates thresholds for detecting changes in the task-relevant variable. As with other threshold-based methods for approaching the stability of suprathreshold perceptual judgments (see Introduction), the extent to which the results may be used to predict the stability such judgments across changes in other scene variables is not known. Experiments that explore this link, perhaps by directly comparing results from the two paradigms with similar stimuli and the same set of observers, are of considerable interest. The results of such experiments might also be helpful in pointing the way to theory that would link results across the two paradigms; at present we do not have such theory in hand (but see Abrams, Hillis, & Brainard, 2007).

Previous authors have suggested that lightness constancy improves with increasing background “articulation”. That is, increasing the number of objects in the background and/or the degree to which their reflectance varies tends to improve constancy (Gilchrist, 2006; Radonjić & Gilchrist, 2013; see also Radonjić, Cottaris, & Brainard, 2015; Kraft, Maloney, & Brainard, 2002). This may on the surface seem in contradiction to our results; we find increasing the variance of the background reflectances has a deleterious effect on lightness discrimination performance. Note, however, that articulation is thought to improve constancy when the task-irrelevant variation is a change in illumination, and where the background itself is held fixed across this change. In our experiments, the illumination is held fixed and we consider the effect of the background per se, with the background change occurring across the two intervals of each forced-choice trial. Thus, we are studying a different aspect of lightness constancy than where increased articulation is thought to lead to improvements, and our results are not in conflict with previous findings.

Our paradigm could be used to study constancy across changes in illumination, if the task-irrelevant variation used in the experiment were in the illumination rather than the background object reflectances. In that case, the articulation idea would predict a smaller elevation of lightness discrimination thresholds when the effect of illumination variation was studied for scenes with higher variance in the background reflectance, as long as the background was held fixed across the two intervals of each trial.

### Applications to Understanding Neural Mechanisms

A longstanding goal of vision science is to connect psychophysical performance to its underlying neural mechanisms. For probing mechanisms that mediate perceptual constancies, our paradigm has the attractive feature that there is a well-defined correct answer on each trial, so that for studies with animal subjects it is possible to provide performance-contingent reward. In addition, there are well-worked out methods for predicting psychophysical discrimination performance from recordings of the responses of neural populations (Shadlen, Britten, Newsome, & Movshon, 1996; Parker & Newsome, 1998; Cohen & Newsome, 2009; Nienborg, Cohen, & Cumming, 2012; Ruff, Ni, & Cohen, 2018), and the theoretical links between such analysis and performance should continue to hold when task-irrelevant stimulus variation is added to the paradigm. Complementing neural measurements that include random, unpredictable task-irrelevant stimulus variation with such analyses may provide rigorous quantitative insights about the sensory-perceptual processing and the neural computations underlying color and lightness constancy specifically, and perceptual constancy more generally.

### Model of Natural Surface Reflectances

We used a truncated multivariate normal distribution as the statistical model for the projection weights of a linear model of naturally occurring reflectances, to sample the background object reflectance functions. This model was developed in our earlier work and is evaluated more fully there (Singh, Cottaris, Heasly, Brainard, & Burge, 2018; see also Brainard & Freeman, 1997; Zhang & Brainard, 2004). The model is based on measurements of the surface reflectance functions of the Munsell papers (Kelly, Gibson, & Nickerson, 1943) as well as natural surfaces characterized by Vrhel (1994). The underlying multivariate normal provides a convenient way to capture two basic aspects of natural variation in reflectance. First, these reflectances are well-described by low-dimensional linear models (Cohen, 1964; Maloney, 1986; Parkkinen, Hallikainen, & Jaaskelainen, 1989). Second, within the reflectance subspace defined by the linear models, not all reflectances are equally likely to occur. Still, we think it likely that future work will lead to more accurate statistical models of naturally occurring reflectance. For example, it is possible that replacing the linear model approach with a prior that favors spectrally-smooth reflectance functions (Jiang, Farrell, & Wandell, 2016) would lead to a more accurate characterization. In addition, we have assumed that the distribution of reflectance functions over objects is independent, but this assumption may not be accurate. Approaches to modeling a dependency have been suggested (Shen & Yeo, 2011; Gehler, Rother, Kiefel, Zhang, & Schölkopf, 2011; Barron & Malik, 2012b; Barron & Malik, 2012a).

It is important to note that the quantitative relation we measured between the magnitude of internal noise and the effect of external noise introduced as variation in background object reflectances depends on how the distribution of naturally-occurring reflectances is modeled. If the model of reflectances overestimates the natural variation, the effect of external noise in natural scenes will be less than we estimated. Conversely, if the model of reflectances underestimates the natural variation, the effect of external noise in natural scenes will be greater than we estimated. Importantly, improved future characterization of naturally occurring reflectances, obtained through the acquisition of additional reflectance measurements and advances in their statistical description, could be used in conjunction with the parameters of the LINRF model formulation, without need for new data collection, to update the estimate of the effect of naturally occurring background object reflectance variation on object lightness perception.

### Rule of Combination

In the present work, we considered variation in only a single task-irrelevant variable. In natural scenes, there are many task-irrelevant variables. In the case of judging object lightness, these include object-extrinsic factors such as the scene illumination, the position and 3D orientation of the target object in the scene, the viewpoint from which the object is viewed, and various object-intrinsic factors like its shape and size. Variation in each of the factors could in principle elevate thresholds for discriminating object lightness. Our paradigm allows characterization of the effect of these task-irrelevant variables and quantifies that effect for each such variable in the same internal-noise referred units. One potentially important future direction is to measure the combined effect of simultaneous variation of multiple task-irrelevant variables, and to test hypotheses about rules of combination that predict the joint effects of such simultaneous variation.

## 6 ACKNOWLEDGEMENTS

NSF BCS 2054900 (VS), NIH RO1 EY10016 (DHB), NIH R01 EY028571 (JB).

## APPENDIX Measurement of object lightness discrimination thresholds under variation in background object reflectances

The control experiment, preregistered as Experiment 2, provided preliminary data that helped shape the design of the main experiment presented in the paper (which was Experiment 3 of the preregistration documents). It aimed to determine whether variation in the reflectance of background objects had an effect on human lightness discrimination thresholds. It established that human object lightness discrimination thresholds increase if the reflectances of background objects vary, as compared to the case when the discrimination is made against a constant background. It also studied the effect of inclusion or not of secondary reflections in the rendering process and assessed the effect of implementing background object reflectance variation across trials rather than across intervals.

The basic methods were the same as for preregistered Experiment 3. The practice session was conducted with the images in Condition 1 described below. The observers were retained for the experiment if their average threshold of the last two blocks during the practice session was lower than 0.030. This was a deviation from the preregistered plan where we set the threshold criterion as 0.025. After collecting data from 8 observers, we realized that the criterion was too strict. Only one observer had met the criterion. After modifying the threshold criterion, we included two of the initially discontinued observers in our experiment (Observer 5 and Observer 8). A total of 11 naïve observers participated in the practice sessions. Four of these observers met the criteria for continuing the experiment. Two of these observers also participated in the main experiment (Observer 4 and Observer 8). The visual acuities of these 4 observers were: Observer 4, L = 20/15, R = 20/20; Observer 5, L = 20/20, R = 20/40; Observer 8, L = 20/30, R = 20/25; Observer 11, L = 20/25, R = 20/30. Observers 5, 8, and 11 wore personal corrective eyewear both during vision testing and during the experiments. Observer 4 did not require or use corrective eyewear.

We measured lightness discrimination threshold of four naïve human observers using a two-interval forced choice paradigm. The thresholds were measured for three specific types of background variation (Figure S1). The reflectance spectra of the background objects were generated with the covariance scalar set to 1. These three conditions were:

### Condition 1. Fixed background

In this condition, the spectra of objects in the background were kept fixed for all trials and for all intervals. We generated 11 images, one at each comparison LRF level.

### Condition 2. Between-trial background variation

In this condition, the spectra of the objects in the background were the same for the two intervals within a trial but varied from trial-to-trial.

### Condition 3. Within-trial background variation

In this condition, the spectra of the objects in the background varied between trials as well as between the two intervals of a trial. The background variation corresponded to covariance scalar equal to 1.

In Conditions 2 and 3, the light reflected from the target object varied from image to image (even at the same LRF level of the target object) because of secondary reflection of light coming from the background objects was included in the rendering. We also measured the thresholds without secondary reflections for these two conditions. We call these conditions Condition 2a and 3a.

**Figure S1:**
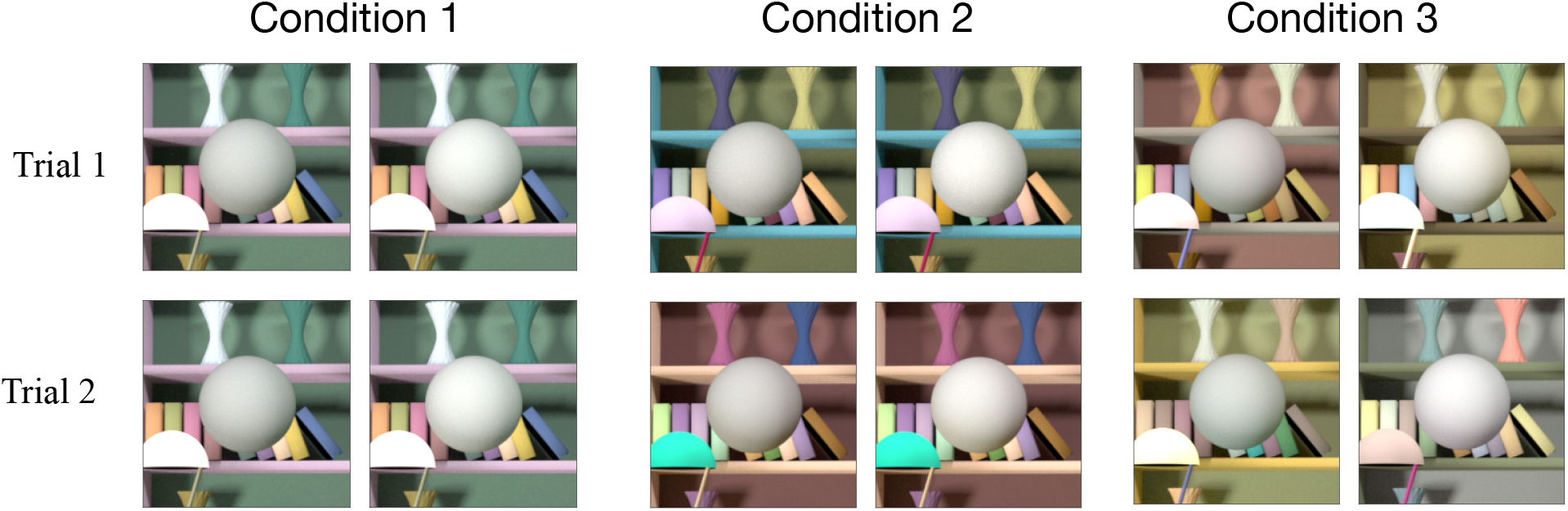
Control experiment stimuli. Example stimuli for Conditions 1, 2 and 3 in the control experiment (preregistered Experiment 2) to study the effect of variation in background object reflectances on lightness discrimination threshold. In condition 1, the background was fixed in every trial and every interval. In Condition 2, the background object reflectances varied from trial to trial, but remained fixed in the two intervals of a trial. In Condition 3, the background object reflectances varied in each trial and interval. For illustration, in this figure we have chosen the stimulus on the left to be the standard image with target object at 0.4 LRF and the on the right to be comparison image with target object at 0.45 LRF. In the experiment, the two images were presented sequentially in random order at the center of the screen. Conditions 2a and 3a stimuli are similar to Conditions 2 and 3 respectively, but without secondary reflections.

### Condition 2a. Between-trial background variation without secondary reflection

Same as Condition 2, but without multiple reflections of light from object surfaces. The light rays only bounce off once from the surfaces before coming to the camera.

### Condition 3a. Within-trial background variation without secondary reflections

Same as Condition 3, but without multiple reflections of light from object surfaces. Condition 3a was the same as the experiment reported in the main paper for covariance scalar equal to 1.

Figure S2 shows the discrimination thresholds of the four human observers for the five conditions studied in this experiment. We plot the mean threshold and the standard error of the mean (SEM) taken over the three separate threshold measurements. For each observer, the thresholds for Conditions 3 and 3a were higher compared to Conditions 1, 2 and 2a. The average increases in threshold of the observers for Conditions 3 and 3a as compared to Condition 1 (baseline) were 79% and 60% respectively. The average increases in threshold for Conditions 2 and 2a were much smaller, 13% and 17% respectively. The thresholds for Conditions 1, 2 and 2a were nearly within one SEM of each other (averaged over the observers and three conditions). On the other hand, the thresholds for Conditions 3 and 3a were respectively (on average) 7.2 and 5.4 SEM larger than the threshold of Condition 1. The thresholds without secondary reflections (Conditions 2a and 3a) were within one SEM from the conditions with secondary reflections (Conditions 2 and 3).

The control experiment established that lightness discrimination thresholds are higher for the case when the two objects are being discriminated against different backgrounds on the same trial, as compared to when the backgrounds are the same within trial. Trial-to-trial variability in background object reflectances across trials has little, if any, effect. The effect is similar when the rendering is performed with and without secondary reflections, indicating the effect is due to the spectral change in the background and not due to the variation in the amount of light being reflected from the target object. In the main experiment, we rendered without secondary reflections to avoid introducing such variability. Figure S2 also shows the threshold of the observers in the main experiment (preregistered Experiment 3) for the condition with covariance scalar equal to 1. This condition is equivalent to Condition 3a of the control experiment (preregistered Experiment 2). Thresholds were consistent across the two measurements.

**Figure S2:**
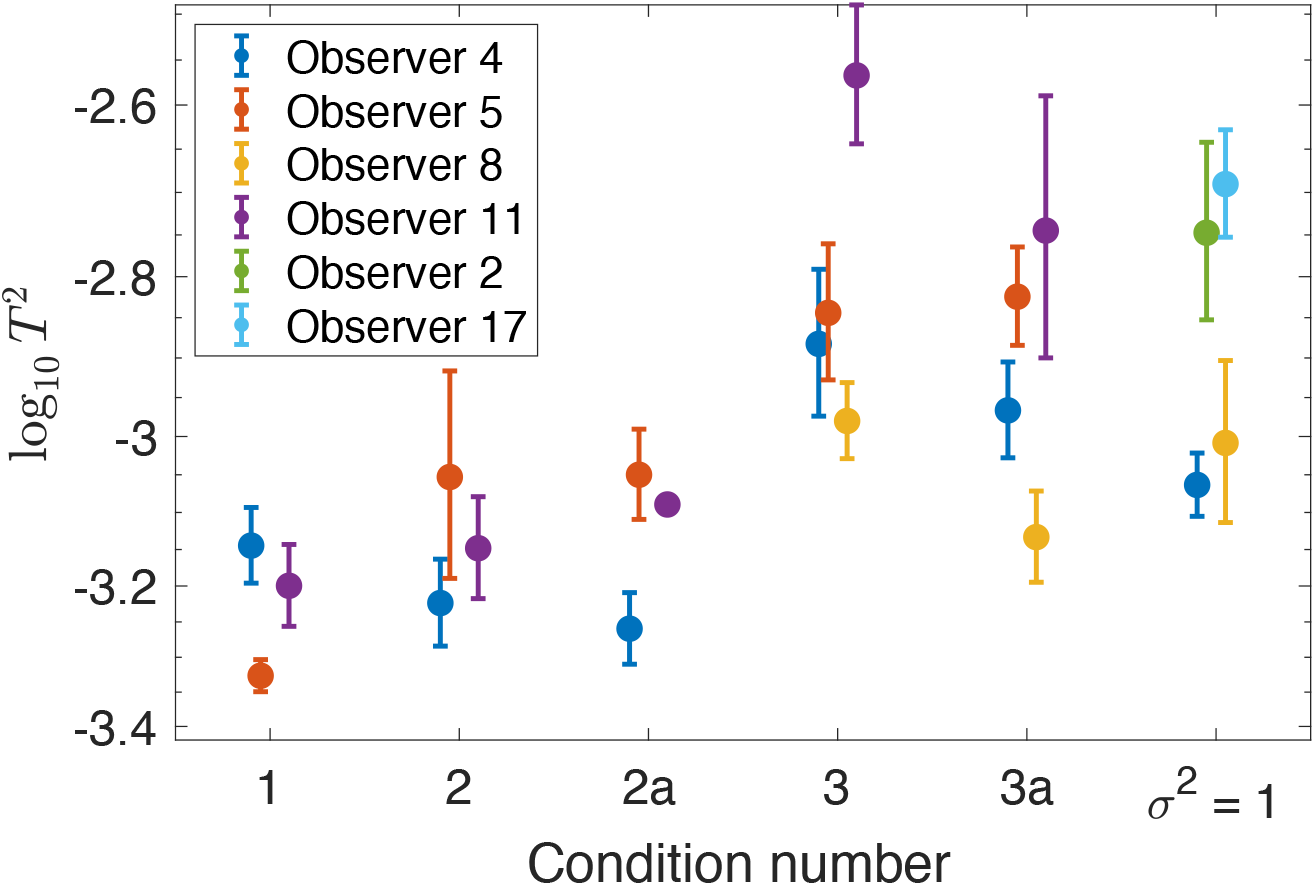
Control experiment. Lightness discrimination threshold of four human observers in the five conditions in the control experiment (preregistered Experiment 2). The plotted points have been jittered horizontally to avoid marker overlap. The thresholds are higher for the condition where the target objects are compared against a change in background object reflectances (Conditions 3 and 3a) than when the background is held fixed within each trial (Conditions 1, 2, 2a). Secondary reflections do not have any significant effect on thresholds (Conditions 2a and 3a). Condition 3a of the control experiment is equivalent to the condition of the main experiment (preregistered Experiment 3) with covariance scalar equal to 1. The thresholds for this condition of the main experiment are plotted here for comparison (*σ*^2^ = 1). Two observers from the control experiment also participated in the main experiment.

**Figure S3:**
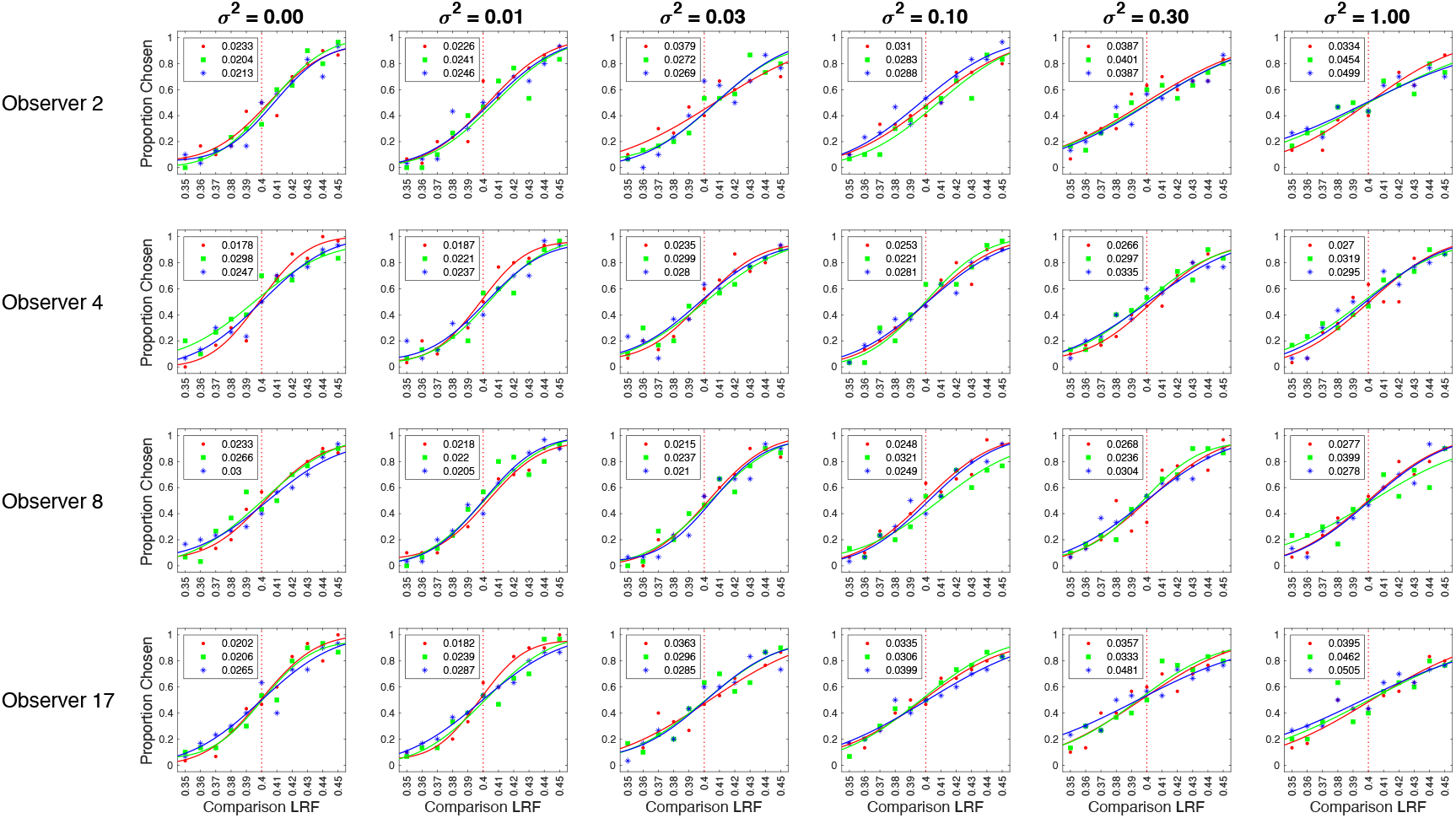
Psychometric functions for all observers. Same as Figure 4 for all observers retained in the main experiment.

**Table S1:** Thresholds for Control Experiment (Preregistered Experiment 2): Mean threshold (averaged over blocks) ± SEM of four human observers for five background variation conditions studied in experiment 2.

| | Mean Threshold ± SEM (averaged over sessions) | | | | |
| --- | --- | --- | --- | --- | --- |
| Observer | Condition 1 | Condition 2 | Condition 2a | Condition 3 | Condition 3a |
| 4 | 0.0269±0.0013 | 0.0254±0.0013 | 0.0235±0.0011 | 0.0366±0.0030 | 0.0330±0.0018 |
| 5 | 0.0217±0.0005 | 0.0305±0.0039 | 0.0300±0.0017 | 0.0382±0.0031 | 0.0389±0.0022 |
| 8 | 0.0167±0.0011 | 0.0169±0.0020 | 0.0175±0.0017 | 0.0325±0.0016 | 0.0273±0.0016 |
| 11 | 0.0252±0.0013 | 0.0268±0.0018 | 0.0285±0.0002 | 0.0525±0.0038 | 0.0439±0.0068 |

**Table S2.** Thresholds for Main Experiment (Preregistered Experiment 3): Mean threshold (averaged over blocks) ± SEM of four human observers measured at six logarithmically spaced values of the covariance scalar.

| Observer | Covariance Scalar |  |  |  |  |  |
| --- | --- | --- | --- | --- | --- | --- |
|  | 0 | 0.01 | 0.03 | 0.1 | 0.3 | 1 |
| 2 | 0.0217±0.0009 | 0.0238±0.0006 | 0.0307±0.0036 | 0.0294±0.0008 | 0.0392±0.0005 | 0.0429±0.0049 |
| 4 | 0.0241±0.0035 | 0.0215±0.0015 | 0.0271±0.0019 | 0.0246±0.0018 | 0.0299±0.0020 | 0.0295±0.0014 |
| 8 | 0.0266±0.0019 | 0.0214±0.0005 | 0.0221±0.0008 | 0.0273±0.0024 | 0.0269±0.0020 | 0.0318±0.0041 |
| 17 | 0.0224±0.0020 | 0.0236±0.0030 | 0.0315±0.0024 | 0.0347±0.0027 | 0.0390±0.0046 | 0.0454±0.0032 |

## Footnotes

1 This type of experiment may be instrumented with instructions that prompt the observer to report how the object appears, or with instructions that prompt the subject to report their estimate of some aspect of the object’s reflectance. Exactly what observers report under either of these instructional regimes, as well as the nature of instructional effects, is an important but thorny issue that we will not digress on further in this paper. See Radonjić and Brainard (2016) for a recent treatment of the issue, as well as the references therein.

2 We adopt the lightness discrimination threshold terminology based on the underlying assumption that observers perform the task using their perceptual lightness representation, and indeed our instructions to subjects used the lightness terminology to describe what should be judged. The actual stimulus variable being varied, however, was the simulated achromatic reflectance of the target object being judged, and feedback was given based on the value of this reflectance. In this paper, we do not explore the question as to whether the results would be affected if we had varied the instructions given to subjects (see footnote 1 above).

3 We use LRF rather than the more generic term albedo as our single number summary of the underlying spectral surface reflectance function, as the LRF is explicit about how variation in reflectance over wavelength should be taken into account.

4 The preregistration documents relevant to this paper are those for Experiments 1, 2 and 3. The site also contains preregistrations for subsequent work not reported in this paper.

5 Here we neglect the effect of the fact that we truncated the distribution to enforce a requirement that reflectance at each wavelength lies between 0 and 1. We return to account for this in the LINRF formulation below.

